# Tumor-intrinsic MHC-II activation in pancreatic ductal adenocarcinoma enhances immune response and treatment efficacy

**DOI:** 10.1101/2025.10.21.683745

**Authors:** Canping Chen, Kyle P. Gribbin, Xi Li, Tugba Y. Ozmen, Furkan Ozmen, Shamilene Sivagnanam, James Kim, Katie E. Blise, Xinxing Yang, Yi Zhang, Roselyn S. Dai, Dove Keith, Mara H. Sherman, Mu-Shui Dai, Lisa M. Coussens, Charles D. Lopez, Rosalie C. Sears, Gordon B. Mills, Katelyn T. Byrne, Zheng Xia

**Affiliations:** Department of Biomedical Engineering, Oregon Health & Science University, Portland, OR, USA; Department of Cell, Developmental and Cancer Biology, Oregon Health & Science University, Portland, OR, USA; Department of Obstetrics and Gynecology, Tongji Hospital, Tongji Medical College, Huazhong University of Science and Technology, Wuhan, China; Division of Oncological Sciences, Knight Cancer Institute, Oregon Health & Science University, Portland, OR, USA; Brenden-Colson Center for Pancreatic Care, Oregon Health & Science University, Portland, OR, USA; Department of Molecular and Medical Genetics, Oregon Health & Science University, Portland, OR, USA; Cancer Biology & Genetics Program, Memorial Sloan Kettering Cancer Center, New York, New York, USA; Knight Cancer Institute, Oregon Health & Science University, Portland, OR, USA; Department of Medicine, Division of Hematology and Medical Oncology, Oregon Health & Science University, Portland, OR, USA; Center for Biomedical Data Science, Oregon Health & Science University, Portland, OR, USA

**Keywords:** Pancreatic cancer, Single-cell sequencing, Spatial transcriptomics, Immunomodulatory therapies, Bioinformatics, Tumor heterogeneity

## Abstract

Pancreatic ductal adenocarcinoma (PDAC) is characterized by an immunosuppressive tumor microenvironment (TME) and poor prognosis. While major histocompatibility complex class II (MHC-II) expression is traditionally associated with professional antigen-presenting cells, its role in PDAC malignant cells remains underexplored. Herein, we utilized single-cell RNA sequencing (scRNA-seq), spatial transcriptomics, bulk RNA sequencing, multiplex immunohistochemistry (mIHC) and ex vivo studies in culture with both human and murine models to investigate the prognostic relevance of MHC-II expression in malignant PDAC cells. Elevated MHC-II expression in malignant cells was strongly associated with increased infiltration of CD4^+^ T and CD8^+^ T cells in human PDAC, and pronounced co-localization with plasma cells, indicative of an antigen-activated immune microenvironment. In the KPC mouse model of PDAC, pharmacologic induction of MHC-II expression by cobimetinib treatment in malignant epithelial cells significantly enhanced the therapeutic response to immune checkpoint blockade (ICB). These findings highlight the role of malignant cell-intrinsic MHC-II expression in promoting antigen presentation and fostering an anti-tumor immune microenvironment. Our results position MHC-II as a promising prognostic biomarker and therapeutic target in PDAC, paving the way for novel immunomodulatory strategies.

**Summary of highlights:** Single-cell and spatial transcriptomic analyses reveal that elevated MHC-II expression in malignant PDAC cells correlates with increased infiltration of CD4⁺ and CD8⁺ T cells.

Stimulating MHC-II expression in tumors effectively enhances immunotherapeutic responses to ICB in the PDAC KPC mouse model, including PDAC tumors previously resistant to therapeutic interventions.

MHC-II serves as a prognostic biomarker and a promising target for immunotherapy in PDAC.

## Introduction

Pancreatic ductal adenocarcinoma (PDAC) has the poorest survival rates among cancers for all stages combined, marked by a median survival of six months and a five-year survival rate < 10%^1^. This aggressive malignancy is characterized by extensive local invasion, distant metastases, and a highly heterogeneous tumor microenvironment (TME) associated with early metastases, rendering most cases ineligible for surgical resection and significantly complicates treatment strategies^2,3^. Moreover, the lack of effective treatment options leads to reliance on cytotoxic therapeutic regimens that fail to achieve significantly improved treatment outcomes^4–6^.

Traditional genomic and transcriptomic analyses have historically relied on bulk tumor data, often neglecting the crucial role of the TME in PDAC progression and immune evasion^7,8^. Additionally, PDAC tumors exhibit marked heterogeneity, characterized by diverse cellular clones and distinct immune landscapes that critically influence tumor behavior and therapeutic resistance^9^. However, recent advancements in single-cell RNA sequencing (scRNA-seq) have provided unprecedented insights into this heterogeneity, enabling a more detailed exploration of the interactions between malignant PDAC cells and the immune microenvironment thus providing a granular understanding essential for refining therapeutic strategies^10–13^.

PDAC tumors are highly complex and includes not only malignant cells, which evolve, giving rise to distinct clones with diverse characteristics, but also a wide variety of stromal and immune cell populations^14^. On one hand, there are cell types with established roles in promoting tumor cell survival and evasion from anti-tumor immune cells, such as cancer- associated fibroblasts (CAFs), tumor-associated macrophages (TAMs), and regulatory T cells (Tregs)^15^. On the other hand, high densities of tumor-killing immune cells, such as CD8⁺ cytotoxic T cells (albeit relatively uncommon in PDAC^16–18^) and CD4^+^ T cells, are associated with improved survival, reflecting a more robust anti-tumor immune response, which has emerged as a critical prognostic factor in PDAC^19–21^.

Despite these insights, the influence of tumor cell-intrinsic factors, particularly major histocompatibility complex class II (MHC-II) expression, on modulating immune responses has been insufficiently explored in PDAC. MHC-II molecules, traditionally expressed on professional antigen-presenting cells (APCs), such as dendritic cells, B cells, and macrophages, present processed exogenous antigenic peptides to CD4^+^ T cells, thereby promoting activation and differentiation of several leukocyte subsets, some of which (such as T cells) can orchestrate adaptive immune responses^22^. Notably, emerging evidence indicates that some malignant cells can express MHC-II, potentially enhancing their immunogenicity and altering the interactions of malignant cells with the immune TME^23,24^.

To address the functional significance of malignant cell expression of MHC-II, we employed a comprehensive multi-omics approach, integrating single-cell and spatial transcriptomic data from both human and mouse models of PDAC, to investigate the role of MHC-II expression. Specifically, we examined how PDAC cell-intrinsic MHC-II expression influenced leukocyte activation and therapeutic responses. Our findings demonstrated that elevated MHC-II expression in malignant PDAC cells correlates with increased infiltration of CD4⁺ and CD8⁺ T cells in the TME, reflecting a more cytotoxic permissive immune microenvironment, which we validated by leveraging a high-resolution single-cell atlas comprising over a million cells across different stages of PDAC progression. Additionally, mechanistic experiments employing the PDAC (KPC) mouse models revealed that induction of MHC-II expression in malignant PDAC cells enabled a shift from checkpoint inhibitor resistance to therapeutic responsiveness. Notably, we observed that the same KPC models, including those which previously showed resistance to combination chemo- and immunotherapy (gemcitabine, nab-paclitaxel, anti-CD40 agonist, anti-CTLA-4, and anti-PD- 1) as reported in Li et al.^25^, exhibited a marked therapeutic response when MHC-II expression was pharmacologically induced via cobimetinib stimulation and subsequently treated with dual PD-1 and CTLA-4 blockade, highlighting the potential of MHC-II pathway activation to sensitize PDAC tumors to ICB.

Collectively, our findings identify tumor cell expression of MHC-II as both a potential prognostic biomarker and a critical modulator of the immune landscape in the pancreatic cancer TME. Therapeutic strategies that enhance MHC-II expression in malignant PDAC cells may represent a novel approach to sensitize the PDAC TME to immunotherapy by boosting malignant PDAC-intrinsic antigen presentation, thereby overcoming resistance and improving clinical treatment outcomes in this highly lethal malignancy.

## Results

### Study design and overview of the study cohort

Given the hypothesis MHC-II expression on malignant cells regulates immune contexture within the TME, we aimed to define its biological and clinical significance using a multi-omics framework. To begin, we analyzed paired pre- and post-treatment PDAC samples from a patient enrolled in a window-of-opportunity clinical trial of the MEK inhibitor cobimetinib (NCT04005690), where treatment led to reduced malignant cell proliferation (decreased Ki- 67) and modulation of MAPK signaling, correlating with a therapeutic response (**Fig. 1A**). To molecularly confirm our observation from this patient, we integrated a range of datasets encompassing 264 patient PDAC specimens, including 195 1.0 mm^2^ tumor regions evaluated by multiplex immunohistochemistry (mIHC) reflecting PDAC samples obtained from 11 patients, 39 murine PDAC RNA-seq samples, and multiple spatial transcriptomics datasets including two Xenium samples, nine Slide-seq samples (ST), and 73 Visium slides. For additional insights into PDAC biology and immune interactions, we included 22 publicly available PDAC scRNA-seq datasets, comprising 1,042,407 cells. Moreover, to validate the findings, data from the Cancer Cell Line Encyclopedia^26^ (CCLE) database with 1,165 cancer cell lines (including 49 PDAC cell lines) were incorporated into the analysis. These comprehensive datasets enabled the integration of clinical, experimental, and computational analyses to decipher how neoplastic cell expression of MHC-II expression correlates with TME and leukocyte features in PDAC **(Fig. 1A)**.

**Figure 1.**
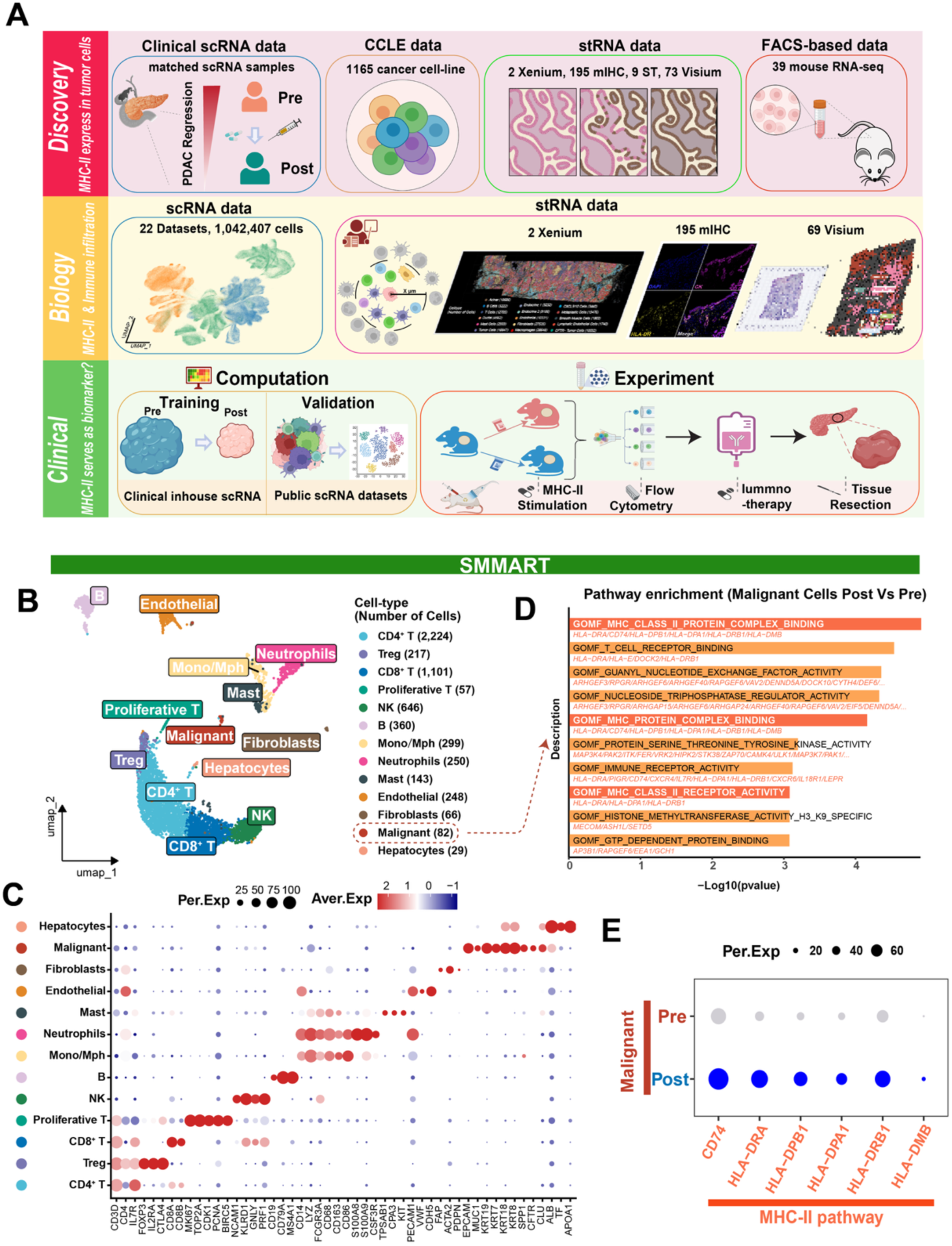
Properties of the cellular TME of pancreatic cancer metastases to the liver. **A.** Study design and datasets overview. **B.** UMAP plot of pre- and on-treatment cell populations with distinct cell types annotated: cells were classified into epithelial (malignant) cells and hepatocytes, stromal cells (fibroblasts, endothelial cells), and immune cell types (CD4⁺ and CD8⁺ T cells, Tregs, natural killer (NK) cells, monocytes/macrophages (Mono/Mph), B cells, mast cells and neutrophils). Each cluster is annotated based on specific marker genes, with the number of cells per cell type indicated. **C.** Dot plot showing average expression levels and percent expression of key marker genes across different cell populations: the dot size represents the percentage of cells expressing a specific gene within a given cell type, while color intensity indicates the average expression level. **D.** Pathway enrichment analysis of on- vs pre-treatment malignant PDAC cells: ClusterProfiler analysis of differentially expressed genes showed significant enrichment of MHC-II-related pathways, particularly those involved in antigen processing and presentation. **E.** Expression of MHC-II pathway genes pre- and on-treatment: dot plot showing key MHC-II-related genes (*CD74*, *HLA-DRA*, *HLA-DPB1*, *HLA-DPA1*, *HLA-DRB1* and *HLA-DMB*) in malignant cells pre- and on-cobimetinib treatment.

### Single-cell RNA analysis of pancreatic ductal adenocarcinoma liver metastases

First, to evaluate transcriptomic reprogramming following MEK1/2 inhibition with cobimetinib, we conducted scRNA-seq on paired pre-treatment and on-treatment PDAC metastatic liver specimens biopsied from a patient (NCT04005690) who received 10 days of treatment with cobimetinib. Using the 10x Genomics platform, we generated high-quality single-cell transcriptomic profiles from a total of 6,187 cells, including 1,734 cells from the pre-treatment sample and 4,453 cells from the on-treatment biopsy sample. Notably, this patient, whose tumor molecularly responded to treatment as evidenced by a significant decrease of malignant cell Ki67 expression and increased infiltration of diverse leukocytes following cobimetinib therapy, provided an opportunity to investigate the underlying mechanisms of a tumor’s response to therapy.

To classify cellular heterogeneity in the metastatic PDAC liver, we combined pre- and on-treatment scRNAseq data (Materials and Methods), normalized to count for sequencing depth, and to mitigate technical variations, and then applied principal component analysis (PCA). Potential batch effects were detected by using Uniform Manifold Approximation and Projection (UMAP), which demonstrated minimal batch effects, as most clusters contained cells from both pre- and on-treatment conditions indicating the tumor of origin was the dominant determinant in the UMAP **(Supplementary** Fig. 1A).

To investigate specific cell subsets, cell clusters were annotated based on expression of well-established cell-type lineage biomarkers^27,28^. Primary cell types identified in the PDAC TME included epithelial cells, stromal cells, and immune cells. Secondary clustering further refined these classifications, revealing key subpopulations including CD4⁺ T cells (*CD3D*, *CD3E*, *IL7R*), regulatory T cells (Tregs) (*FOXP3*, *IL2RA*, *CTLA4*), CD8⁺ T cells (*CD8A*, *CD8B*), proliferative T cells (*MKI67*, *TOP2A*, *CDK1*, *PCNA*, *BIRC5*), NK cells (*NCAM1*, *KLRD1*, *GNLY*, *PRF1*), B cells (*CD19*, *CD79A*, *MS4A1*), myelomonocytic cells, e.g., monocytes or macrophage (*CD14*, *LYZ*, *FCGR3A*, *CD68*, *CD163*, *CD86*), neutrophils (*S100A8*, *S100A9*, *CSF3R*), mast cells (*TPSAB1*, *CPA3*, *KIT*), endothelial cells (*PECAM1*, *VWF*, *CDH5*), fibroblasts (*FAP*, *ACTA2*, *PDPN*), malignant tumor cells (*EPCAM*, *MUC1*, *KRT19*, *KRT7*, *KRT18*, *KRT8*, *SPP1*, *CFTR*, *CLU*) and hepatocytes (*ALB*, *TF*, *APOA1*). (**Fig. 1B, 1C**).

Given our interest in antigen presentation by tumor cells, we wanted to ensure high-confidence identification of malignant PDAC cells in subsequent treatment response analyses. To this end, we assessed chromosomal-scale copy number variations (CNVs) in malignant epithelial cells using the InferCNV program^29^. By comparing each gene expression of each cell across chromosomes to reference diploid cells, we identified somatic CNV changes. Malignant epithelial cells exhibited significant CNVs spanning entire p or q arms **(Supplementary** Fig. 1B), indicating a high degree of aneuploidy, a hallmark of PDAC^30,31^. Importantly, these chromosomal alterations were not observed in other cell types, validating the epithelial cells as malignant and underscoring the chromosomal instability characteristic of this metastatic PDAC.

### Significant alteration in MHC-II pathway activity within malignant cell populations on-therapy

Using the clinical biopsy sample, we next investigated the expression profile change in the PDAC TME. After isolating the malignant cell population, we performed differential gene expression (DEG) analysis to assess treatment-induced functional changes. Pathway enrichment using the enricher function in the R package clusterProfiler^32^ (Materials and Methods) revealed marked upregulation of immune-related pathways, indicating a robust immune activation signature within the tumor microenvironment^33^. Notably, three MHC-II- related GO terms consistently ranked among the top enriched pathways—MHC class II protein complex binding, MHC protein complex binding, and MHC class II receptor activity (**Fig. 1D**). Among these, MHC class II protein complex binding was the most significantly enriched and was selected for further analysis. This was supported by significant upregulation of key MHC-II genes, including *CD74*, *HLA-DRA*, *HLA-DPB1*, *HLA-DPA1*, *HLA- DRB1*, and *HLA-DMB* in malignant PDAC cells from on-treatment compared to pre-treatment biopsies (**Fig. 1E**). These findings highlight the functional plasticity of malignant cells in response to therapy^34^ —a critical factor contributing to intercellular heterogeneity in therapeutic response and immunogenicity, both of which can profoundly influence treatment efficacy and patient prognosis^35,36^.

Upregulation of the MHC-II antigen presentation pathway activity in malignant cells during therapy, accompanied by shifts in immune and malignant cell proportions (**Supplementary** Fig. 1C) and a notable decline in the biomarker CA19-9 in this donor, indicated that PDAC cells may be responsive to cobimetinib treatment, potentially correlating with increased expression of antigen presentation machinery on the tumor cell surface.

### Validation of MHC-II pathway gene expression in malignant tumor cells via multi-omics approaches

Before establishing the MHC-II pathway activity as a viable treatment target in PDAC, it was necessary to confirm that MHC-II was indeed expressed at the protein level in malignant tumor cells. Although our scRNA-seq analysis identified that MHC-II and correlated genes were expressed in PDAC cells using cell-type markers **(Fig. 1C)** and copy number variations **(Supplementary** Fig. 1B), the possibility of signal contamination from non-malignant cells remained a concern.

We next analyzed PDAC malignant cell-intrinsic MHC-II expression using transcriptome profiles from the CCLE dataset, which exclusively containing tumor cell-line data. This approach minimized the likelihood of contamination from immune cell-derived MHC-II signals. Our analysis revealed variable MHC-II expression across multiple cancer types, including both solid and hematologic malignancies. A heatmap of MHC-II gene expression highlighted substantial variability between PDAC cancer cell lines **(Fig. 2A)** as well as across all tumor cell lines **(Supplementary Table 1)**, indicating that differential MHC-II expression may underlie variations in therapeutic responses across tumors.

**Figure 2.**
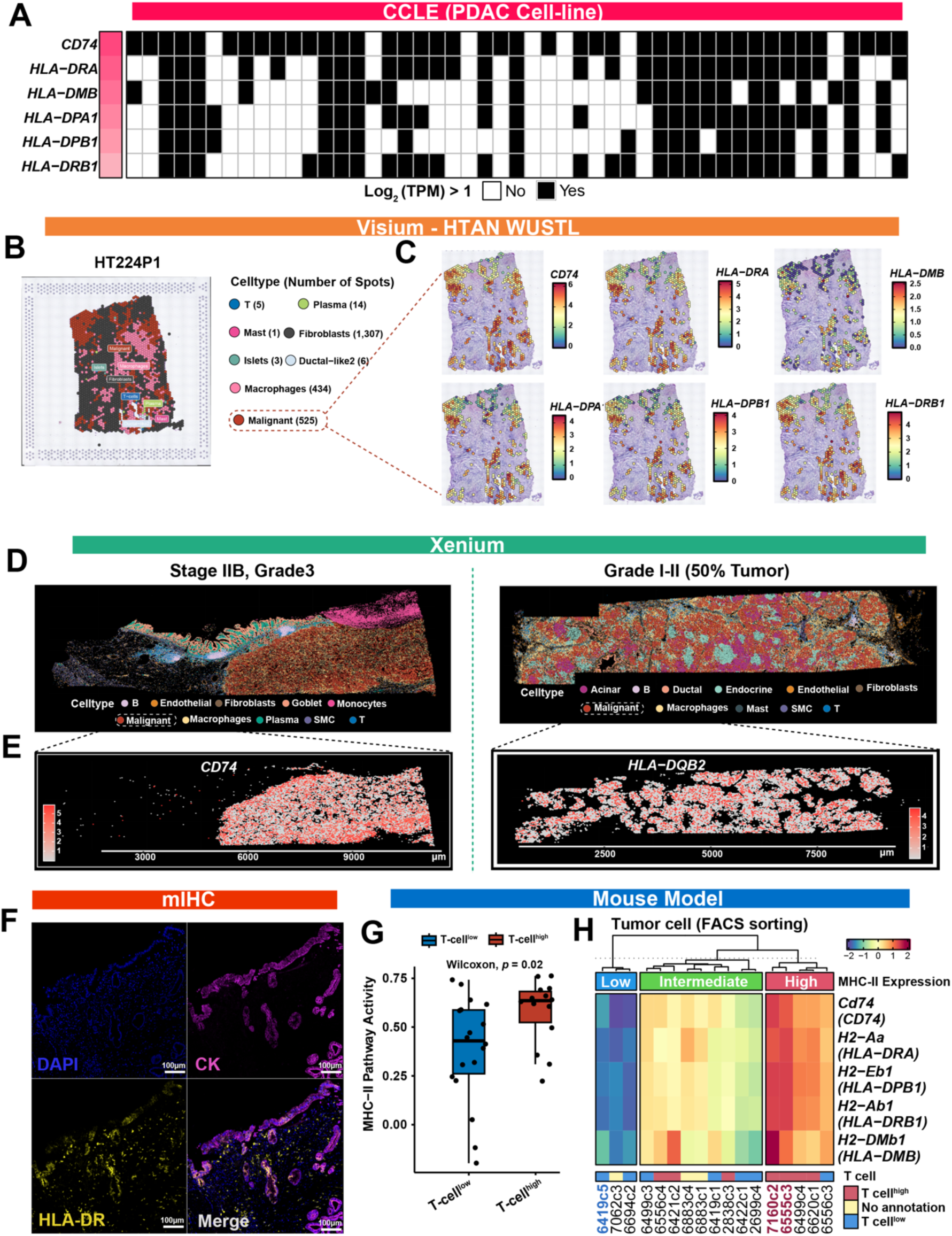
Validation of malignant cell-intrinsic MHC-II expression across multi-omics platforms. **A.** Heatmap of MHC-II gene expression in PDAC cell lines from the CCLE. **B.** Visualization of cell type annotations mapped to Visium spots, inferred by the RDCT deconvolution algorithm. **C.** Spatial distribution of key MHC-II-related genes (*CD74*, *HLA-DRA*, *HLA-DPB1*, *HLA-DPA1*, *HLA-DRB1* and *HLA-DMB*) expression within malignant cell spots in Visium samples, demonstrating variation in MHC-II pathway activation across the tumor microenvironment. **D.** Cell-type resolved spatial maps of two tumor samples profiled by Xenium in situ platform: a Stage IIB, Grade 3 tumor (left) and a Grade I–II tumor with ∼50% malignant content (right). **E.** Spatial gene expression heatmaps of *CD74* (left) and *HLA-DQB2* (right) in malignant regions from the corresponding tumor sections in (**D**). **F.** mIHC analysis of HLA-II expression in PDAC tissues: Representative images showing DAPI (nuclear stain, blue), cytokeratin (CK; epithelial marker, magenta), and HLA-DR (MHC-II marker, yellow) expression. Malignant cells co-staining for HLA-DR and CK indicate MHC-II- positive malignant cells. **G.** Boxplot illustrating comparison of MHC-II pathway activity between T-cell^high^ and T-cell^low^ regions. **H.** Heatmap of key MHC-II-related genes expression in malignant clonal cells sorted via flow cytometry from the KPC mouse model from Li et al^25^. Samples with red or blue color indicate clones used in subsequent studies.

Further validation was performed using 14 Visium spatial transcriptomics samples and matched single-nucleus RNA sequencing (snRNA-seq) data from the HTAN WUSTL atlas^28^ and 55 Visium spatial transcriptomics samples from GSE277783^37^. Using the Robust Cell Type Decomposition (RCTD)^38^ algorithm, we transferred cell-type labels from snRNA-seq or scRNA-seq to spatial transcriptomic spots, deconvolving the cell-type composition at each spot. A representative image is shown in **Fig. 2B**, with additional examples in **Supplementary** Fig. 2. Gene analysis revealed heterogeneous MHC-II expression in malignant cells, as illustrated by representative images in **Fig. 2C**. Pathway enrichment analysis further corroborated these findings, as shown in **Supplementary** Fig. 3. To strengthen these observations, we performed additional validation using three independent spatial transcriptomics PDAC datasets obtained from SORC database^39^, accompanied by corresponding histologic images, which consistently confirmed MHC-II gene expression within malignant cell spots. **(Supplementary** Fig. 4).

To validate the above transcriptomic data, we next utilized two *in situ*^40^ Xenium gene expression datasets (10x Genomics) derived from human PDAC tissue sections of differing grades, using a pre-designed multi-tissue and cancer panel. This allowed for additional validation of MHC-II on tumor cells at the single-cell level. The two datasets encompassed 184,273 cells with 380 genes and 190,962 cells with 474 genes, respectively, captured the expected pancreatic cell types (**Fig. 2D**). By focusing specifically on malignant tumor cell populations, we observed substantial heterogeneity in MHC-II gene expression among cells (**Fig. 2E**), thereby highlighting the variability of MHC-II expression across malignant cell regions and aligning with our scRNA-seq findings.

To further validate MHC-II protein expression on PDAC cells, we performed multiplex immunohistochemistry^41,42^ —a technique that enables simultaneous detection of multiple protein markers within a single tissue section—on resection samples from 11 PDAC patients who had received standard-of-care neoadjuvant chemotherapy, analyzing 195 regions spanning diverse tissue areas and collectively covering 568 mm². Representative images showed DAPI (blue), CK (magenta), and HLA-DR (yellow) staining **(Fig. 2F)**. Quantitative analysis of HLA-II expression confirmed the presence of MHC-II protein in malignant PDAC cells, as illustrated by a representative image in **Supplementary** Fig. 5. Together, these human data provide compelling transcriptomic and proteomic evidence that MHC-II is variably expressed on malignant PDAC cells, supporting the rationale to investigate the tumor-intrinsic regulation of MHC-II in controlled preclinical models.

Given that patient data revealed the PDAC malignant cells express MHC-II in therapy-naïve tumors, with expression further enhanced after cobimetinib treatment (**Fig. 1E**), we therefore sought to delineate the tumor-intrinsic and microenvironmental conditions that induce MHC-II expression in PDAC. To this end, we examined MHC-II gene expression in sorted malignant tumor cells from clonal tumor cell lines derived from KPC mice, as reported by Li et al.^25^, where tumors were categorized into three groups based on immune infiltration. Unsupervised clustering revealed two distinct patterns of MHC-II expression in malignant tumor cells, where elevated MHC-II levels strongly associated with tumors enriched in T cells **(Fig. 2G)**. Our analysis confirmed increased MHC-II pathway activity in tumor cells that gave rise to highly T-cell infiltrated TMEs as compared to tumors that resulted in T cell-low TMEs **(Fig. 2H)**.

These results collectively underscore the heterogeneity of MHC-II expression across PDAC cells in different regions of the TME and the potential connection of tumor cell- expressed MHC-II with immune infiltration to the tumor site.

### Single-Cell atlas of human pancreas

To overcome the limitations of sample size in the previous scRNA-seq study and to provide a comprehensive overview of MHC-II across PDAC progression, we compiled publicly available datasets from the Deeply Integrated Single-Cell Omics (DISCO) database^43^. This effort resulted in creation of a robust Human PDAC Cell Atlas, encompassing over 1 million cells from 264 samples **(Supplementary Table 2)**. The atlas offers a high-resolution view of various stages of PDAC, including normal healthy tissue, normal adjacent tissue, primary PDAC, and metastatic PDAC.

To ensure data consistency across different conditions, we applied batch correction using the BBKNN^44^ algorithm in Scanpy, specifying each research dataset as the batch_key. This approach effectively harmonized data while preserving biological variability. A cohesive temporal perspective of cell populations across PDAC stages was visualized using UMAP, generated with the sc.tl.umap function and default parameters **(Fig. 3A).**

**Figure 3.**
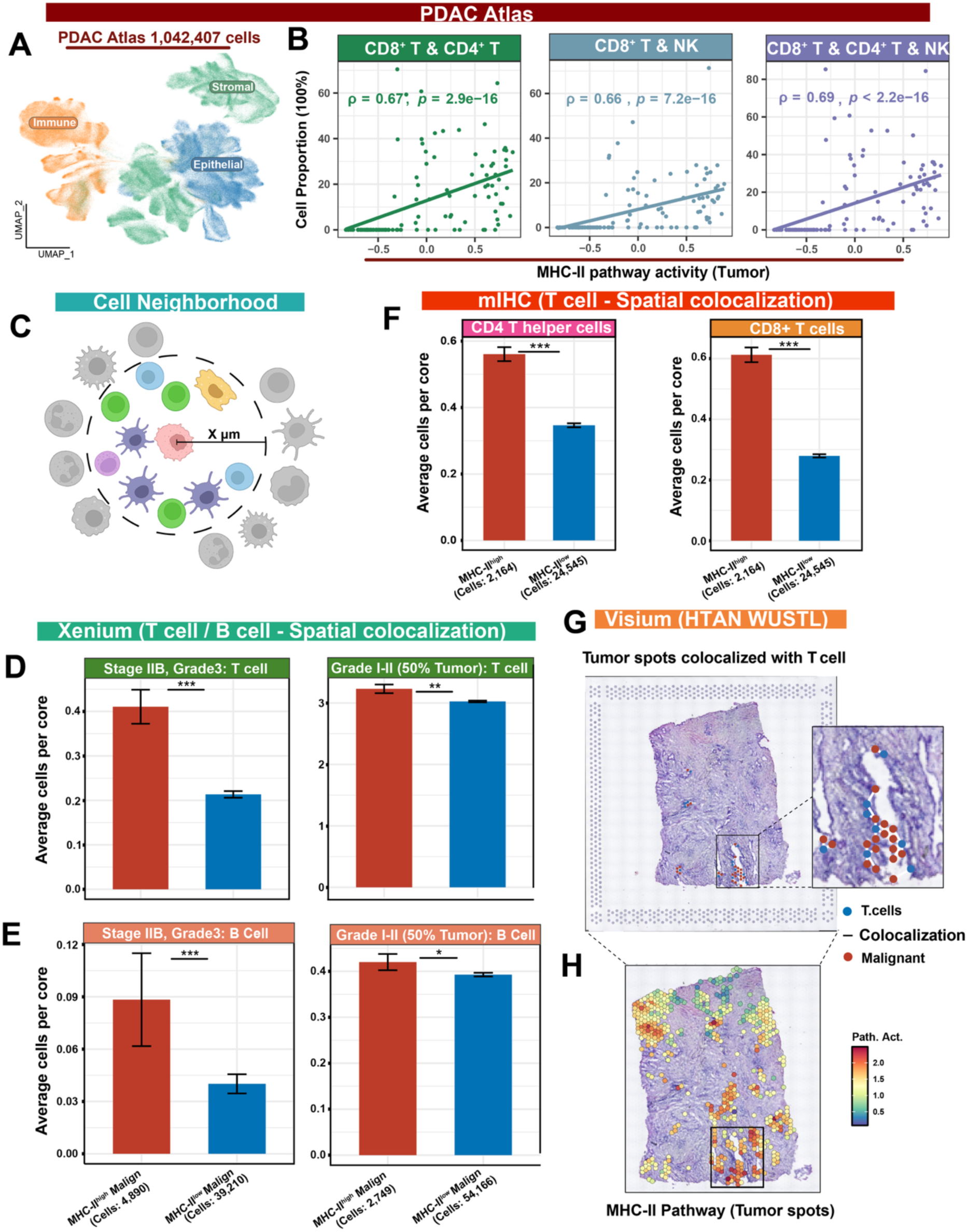
Spatial analysis of MHC-II positive malignant cells and their immune contexture in PDAC. **A.** UMAP of over 1 million single cells from the PDAC Atlas, categorized into epithelial, stromal, and immune compartments. **B.** Correlation between MHC-II pathway activity and immune cell combinations: scatter plots showing positive correlations between MHC-II pathway activity in malignant cells and the proportions of CD8+T cells and CD4+ T cells (left), CD8+ T cells and NK cells (middle), and combined CD8+ T cells, CD4+ T cells, and NK cells (right). **C.** Illustration of spatial neighborhoods around malignant cells: defining a specified radius around each seed cell (orange) to assess the surrounding cellular composition. **D-E.** Quantification of immune cells within malignant cell neighborhoods with differential MHC-II expression (Xenium). Bar plots comparing the average number of T cells **(D)** and B cells (**E**) per core in MHC-II^high^ versus MHC-II^low^ malignant regions across two tumor samples: Stage IIB, Grade 3 (left) and Grade I–II with ∼50% tumor content (right). **F.** Quantification of CD4^+^ T helper cells and CD8^+^ T cells per neighborhood mIHC ROIs, demonstrating significant differences in cell-type abundance between the two groups. **G.** Visium spatial transcriptomics from the HTAN-WUSTL cohort shows malignant cell spots colocalized with T cells are enriched for high MHC-II pathway activity. **H.** Feature-plot of MHC-II pathway activation scores across malignant cell spots further highlights spatial heterogeneity and preferential immune proximity to MHC-II^high^ malignant cell regions. *: *p* < 0.05, **: *p* < 0.01 and ***: *p* < 0.001. The same significance thresholds apply to all subsequent figures unless otherwise specified.

Malignant cells were identified through the R package scATOMIC^45^ (v2.0.2) using the parameter known_cancer_type = "PAAD cell" with default settings for other parameters. This rigorous annotation ensured accurate classification of malignant populations, laying the groundwork for deeper analyses of PDAC progression and heterogeneity.

### High MHC-II pathway activity in malignant cells correlates with enhanced lymphocyte infiltration

In-depth analyses of sorted tumor cells from PDAC tumor-bearing mice revealed that MHC-II pathway activity was significantly elevated in malignant cells from TMEs with high T cell content as compared to TMEs with low T cell content (**Fig. 2G**). To extend this analysis to human PDAC and elucidate a link between MHC-II pathway activity and lymphocyte presence, we conducted a Spearman’s correlation analysis to evaluate the association between MHC-II pathway activity in malignant cells and infiltration levels of various T and natural killer (NK) cell lineages across multiple samples, where malignant cells from each sample were aggregated to create a pseudo-bulk profile for the analysis (Materials and Methods).

The analysis revealed a strong positive correlation between MHC-II activity in malignant cells and the abundance of T cells in the corresponding patient TME (**Fig. 3B**). Notably, combinations of immune cells, such as CD8⁺ and CD4⁺ T cells, and NK cells, and combinations involving CD8⁺ T cells, CD4⁺ T cells, and NK cells, showed particularly high correlations (**Fig. 3B**). Additionally, MHC-II pathway activity in malignant cells was strongly correlated with the proportions of individual lymphocytes and combinations thereof, while showing a negative correlation with endothelial cells and fibroblasts (**Supplementary** Fig. 6A).

Interestingly, despite the overall trend, we identified several samples with high MHC-II pathway activity in the absence of detectable lymphocyte infiltration. Notably, 54 out of 57 such cases originated from the GSE202051 dataset^46^, which was generated using snRNA-seq of frozen tumor samples. Because snRNA-seq is particularly effective at profiling malignant and stromal cells but exhibits reduced sensitivity for capturing lymphocytes^47,48^, this likely accounts for the observed discrepancy. Nonetheless, the strong positive association between MHC-II pathway activity and immune infiltration across multiple independent datasets underscores the robustness of MHC-II pathway activity as a surrogate indicator of immune presence within the tumor microenvironment.

### Lymphocytic spatial neighborhoods reveal distinct spatial landscapes associated with malignant cells exhibiting different MHC-II pathway activity

Tumor cell expression of MHC-II pathway activity has not yet been systematically investigated for its potential role in shaping intratumoral spatial heterogeneity of T cell infiltration. Patients with distinct immune and stromal infiltration patterns often experience markedly different clinical outcomes, particularly in their response to immunotherapy^49,50^. Despite this, current treatment strategies largely overlook the unique composition and spatial organization of tumor-infiltrating immune cells, even though immune contexture, including CD4⁺ and CD8⁺ T cell infiltration, has demonstrated stronger prognostic value than traditional TNM staging^51^.

Given the observed correlation between MHC-II pathway activity in malignant PDAC cells and immune infiltration, we applied the K-nearest neighbor (KNN) algorithm to explore whether malignant cells with high MHC-II level were spatially closer to specific lymphocytic lineages. To do so, we defined “neighborhoods” as physical groupings of cells within a set distance (50 μm) from each malignant cell, referred to as the “seed cell” **(Fig. 3C)** (Materials and Methods).

Within the Xenium spatial transcriptomic datasets, which demonstrated robust T cell infiltration at single-cell resolution, patient malignant tumor cells were stratified into two groups based on MHC-II pathway activity: MHC-II^high^ and MHC-II^low^. Neighborhood composition analysis revealed that T cell abundance was significantly higher in the microenvironments surrounding MHC-II^high^ malignant cells compared to those adjacent to MHC-II^low^ malignant cells **(Fig. 3D)**. This trend remained consistent after correcting for total cell number **(Supplementary** Fig. 6B). A similar enrichment pattern was observed for B cells, with increased representation around MHC-II^high^ malignant cells (**Fig. 3E and Supplementary** Fig. 6C).

The robustness of spatial patterns observed in the Xenium data was next validated through neighborhood analyses at the proteomics levels using mIHC analyses (**Supplementary** Fig. 5 and **Fig. 3F**) and in Visium data sets (**Fig. 3G**), yielding consistent results across multiple omics platforms and different data sets and patient cohorts. Notably, even in tumor slides with only a limited number of T cells detected via Visium spatial transcriptomics, the malignant cells that co-localized with T cells were predominantly those exhibiting the highest levels of MHC-II gene expression within the tumor section **(Fig. 3G - H)**.

Given the established role of plasma cells in modulating immune responses and their potential contribution to antitumor immunity in PDAC^37,52^, we next sought to explore whether spatial proximity to malignant cells with high MHC-II activity might extend beyond T cells to include B cells and specifically, plasma cells. Notably, although the overall proportion of plasma cells did not exhibit a strong correlation with MHC-II pathway activity in malignant cells (**Supplementary** Fig. 6A), CellTrek analysis^53^ further revealed that plasma cell– enriched regions were in significantly closer proximity to MHC-II^high^ malignant cell spots compared to MHC-II^low^ malignant cell spots across all Visium samples that had matched snRNA-seq data (**Supplementary** Fig. 7). This spatial association indicates a preferential interaction between plasma cells and MHC-II^high^ malignant cell regions, emphasizing the spatial heterogeneity of the TME and underscoring the pivotal role of MHC-II pathway activity in orchestrating immune cell dynamics and influencing tumor behavior in PDAC.

### CAF support for hypoxic environments in MHC-II^low^ malignant cell niches

Cancer-associated fibroblasts (CAFs) are one of the most abundant cell populations in the TME and have garnered significant attention for their multifaceted roles in tumor progression^54,55^. Through their interactions with stromal components and immune cells, CAFs drive key processes such as angiogenesis, extracellular matrix (ECM) remodeling, and immune evasion^56^.

Fibroblasts proportion was found to exhibit a negative correlation with MHC-II pathway activity in malignant cells (**Supplementary** Fig. 6A). To further dissect the complex spatial architecture of the TME, we applied nine distinct CAF subtype gene signatures defined in Cords’ research^54^, classifying CAFs based on functional and phenotypic diversity, including vascular CAFs (vCAFs), matrix CAFs (mCAFs), interferon-responsive CAFs (ifnCAFs), tumor-like CAFs (tCAFs), inflammatory CAFs (iCAFs), dividing CAFs (dCAFs), reticular-like CAFs (rCAFs), and antigen-presenting CAFs (apCAFs).

Spatial analysis of these CAF subtypes across primary tumors and metastatic lesions (liver, lung, and peritoneum) revealed distinct patterns of co-localization with malignant cells exhibiting varying MHC-II pathway activity (**Supplementary** Fig. 8). Notably, apCAFs and iCAFs preferentially co-localized with MHC-II^high^ malignant cell spots in the PDAC TME. In particular, apCAF signature scores were consistently elevated in CAF spots adjacent to MHC-II^high^ malignant cell regions across nearly all samples. Given their capacity for antigen presentation, apCAFs may contribute to lymphocyte modulation in the PDAC TME^57^. iCAFs, known for secreting pro-inflammatory cytokines, may similarly support immune modulation in the TME^54^. Conversely, mCAFs and tCAFs were enriched around MHC-II^low^ malignant cell regions. mCAFs specialize in extracellular matrix production and remodeling—processes integral to structural support and tumor progression—while tCAFs were associated with tumor-like phenotypes, metabolic adaptation, and survival signaling.

Importantly, the spatial associations between CAF subtypes and malignant cell MHC-II status were consistent across metastatic sites, sexes, and distinct PDAC tumor subtypes in our analyses (including classical, intermediate, and basal) (**Supplementary** Fig. 8), indicating that these interactions are governed by a complex interplay between intrinsic tumor programs and microenvironmental cues.

While previous studies have reported that malignant cells associated with myCAFs contribute to the exclusion of plasma cells from the TME—a phenomenon linked to poor outcomes and resistance to chemotherapy—these findings have largely been limited to basal-like PDAC tumors^37,58^. In contrast, our results reveal that MHC-II^high^ tumor cells reside within a more immunologically active niche, characterized by increased spatial proximity to various immune cell types, including T cells, B cells and plasma cells, as well as pro-inflammatory and immune-supportive CAF subtypes such as antigen-presenting CAFs and inflammatory CAFs, while exhibiting spatial exclusion from tumor-promoting CAF subsets (myCAFs and tCAFs). These findings indicate that MHC-II pathway activity plays a central role in orchestrating the spatial immune–stromal interactions in the TME and shaping a more permissive immune microenvironment, independent of classical, intermediate or basal-like PDAC subtype classifications.

### Elevated MHC-II expression predicts immunotherapy response

Given the observed association between tumor cell MHC-II activity and lymphocyte infiltration, we next asked whether MHC-II expression could serve as a predictive biomarker for therapeutic responsiveness, particularly in the context of immunotherapy. Thus, we developed a model for predicting MHC-II activity to assess malignant cell responsiveness to drug treatment using the MHC-II pathway signature gene list. The model successfully distinguished sample heterogeneity (**Fig. 4A**) and demonstrated robust predictive performance, as evidenced by its strong correlation with lymphocyte infiltration. Specifically, the percentage of predicted responsive malignant cells showed highly positive associations with combinations of CD8⁺ and CD4⁺ T cells, as well as broader interactions involving CD8⁺ T cells, CD4⁺ T cells, and NK cells (**Fig. 4B**). Detailed correlations for individual cell types and other combinations are provided in **Supplementary** Fig. 9.

**Figure 4.**
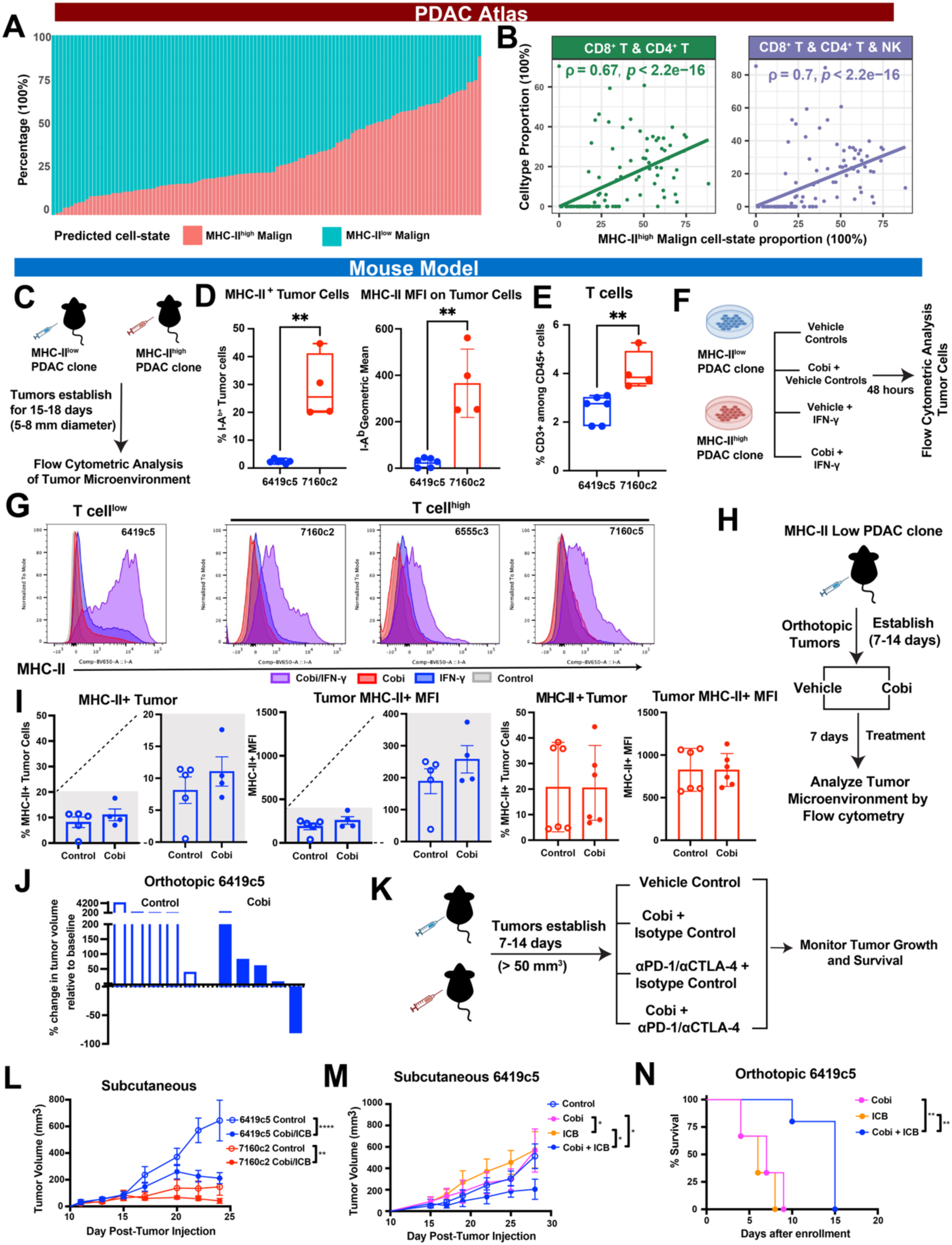
Malignant cell-intrinsic MHC-II expression correlates with immune infiltration and predicts immunotherapy responsiveness. **A.** Single-cell analysis of MHC-II pathway activity in malignant cells. Bar plot depicting predicted response status based on MHC-II expression levels across different PDAC tumor samples. **B.** Correlation of MHC-II^high^ malignant cells with immune cell infiltration: scatter plots illustrating positive correlations between the proportion of MHC-II^high^ malignant cells and immune cell combinations, including left: CD8⁺ and CD4⁺ T cells and right: CD8⁺ T cells, CD4⁺ T cells, and NK cells combined. **C.** Experimental design for in vivo flow cytometric profiling of the tumor microenvironment using syngeneic MHC-II^low^ (6419c5) and MHC-II^high^ (7160c2) PDAC clones. **D.** Percentage of MHC-II⁺ malignant cells (left) and mean fluorescence intensity (MFI) of MHC-II on malignant cells (right) in MHC-II^low^ (6419c5) versus MHC-II^high^ (7160c2) tumor clones. Data are shown as box-and-whisker plots, with boxes indicating the interquartile range, the line marking the median, and whiskers representing the full range. **E.** Frequency of CD3⁺ T cells among CD45⁺ immune cells in 6419c5 and 7160c2 tumor clones. **F.** Schematic of in vitro stimulation with IFN-γ and cobimetinib to induce MHC-II expression, followed by flow cytometric analysis. **G.** MHC-II expression on tumor cells with varying T cell levels following treatment with IFN-γ and cobimetinib in vitro. Histogram overlays showing MHC-II expression in tumor cells categorized as T cell-low (6419c5) and T cell-high (7160c2, 6555c3, 7160c5), also shown above and in right hand corner. Tumor cells were treated with IFN-γ (blue), cobimetinib (Cobi, red), or a combination of both (Cobi/IFN-γ, purple), with untreated control cells shown in gray. Data are normalized to mode to illustrate the shift in MHC-II expression across conditions. **H.** Experimental design for evaluating the impact of cobimetinib on MHC-II expression in orthotopic PDAC tumors. Mice were orthotopically implanted with either MHC-II^low^ or MHC-II^high^ PDAC clones and allowed to establish tumors for 7–14 days. Mice were then treated with either vehicle or cobimetinib for 7 days. **I.** Effect of cobimetinib treatment on MHC-II expression in vivo on malignant clonal cells. Bar graphs showing the percentage of MHC-II+ tumor cells (left) and the MFI of MHC-II (right) on tumor cells in control and cobimetinib-treated tumors from mice bearing indicated tumors. The left-hand side represents data from MHC-II^low^ tumors (6419c5, blue), while the right-hand side represents data from MHC-II^high^ tumors (7160c2, red). Data are displayed as mean ± SEM, with individual data points shown. **J.** Tumor volume changes following cobimetinib treatment: Waterfall plots show the percentage change in tumor volume relative after 14 days of treatment as compared to baseline in individual mice treated with either control or cobimetinib in MHC-II^low^ (6419c5, blue) tumors as shown. Each bar represents an individual mouse. **K.** Experimental design: Mice were subcutaneously implanted with MHC-II^low^ PDAC clones (6419c5 or 7160c2) and allowed to establish tumors (∼50 mm²) over 7–14 days. Mice were then treated with cobimetinib in combination with either dual isotype controls or dual immune checkpoint blockade (ICB; anti-PD-1 and anti-CTLA-4) and monitored for tumor growth. **L.** Tumor volume (mm³) over time in mice bearing 6419c5 or 7160c2 tumors, treated with either control or Cobi/ICB treatment. Data are represented as mean ± SEM. **M.** Tumor growth in subcutaneous 6419c5 tumors. Tumor volume (mm³) over time in mice treated with control, immune checkpoint blockade (ICB; anti-PD-1 and anti-CTLA-4), or a combination treatment of cobimetinib and ICB. Data are presented as mean ± SEM. **N.** Survival analysis in orthotopic 6419c5 MHC-II^low^ tumors (injected and monitored as in **H**), with indicated treatments (as in **M**), with Kaplan-Meier survival curves shown.

To functionally validate the relationship between tumor-intrinsic MHC-II expression and treatment response, we next turned to preclinical mouse models allowing direct experimental manipulation of MHC-II levels in malignant cells. To rigorously assess the impact of MHC-II expression on therapeutic efficacy, both in vitro and vivo experiments were conducted using the orthotopic mouse model with clonal tumor cells expressing varying levels of MHC-II. Tumor cells were specifically engineered to express the YFP lineage tag (KPCY), enabling precise tracking of tumor behavior and treatment responses^25,59^. This model, which harbors the heterozygous *Trp53* mutation (*Trp53^R172H^*), which faithfully recapitulates the histopathological and molecular characteristics of human pancreatic ductal adenocarcinoma.

Flow cytometry (**Fig. 4C**) revealed that tumor cells with high MHC-II activity score from **Fig. 2H** (7160c2) exhibited a significantly higher proportion of MHC-II-positive malignant cells compared to tumor cells with a low MHC-II activity score (6419c5) (**Fig. 4D**).

Correspondingly, tumor cells with elevated MHC-II activity in vivo (**Fig. 4D**) demonstrated significantly increased CD3⁺ T cell infiltration (**Fig. 4E**), supporting the role of MHC-II in fostering an immunologically active tumor site.

Given this positive correlation between MHC-II expression and lymphocyte infiltration, we next explored whether cobimetinib could induce MHC-II expression in MHC-II^low^ tumor cells and potentially augment lymphocyte infiltration and thereby responses to immunotherapy. In vitro, flow cytometry confirmed that tumor cell expression of MHC-II—initially low in all untreated tumor cells—was significantly upregulated following cobimetinib and IFN-γ treatment, especially the MHC-II^low^ tumor cells (**Fig. 4F–G**), indicative of a tumor-cell intrinsic upregulation of MHC-II after cobimetinib treatment.

To evaluate the in vivo effects of cobimetinib, MHC-II expression was assessed on-treatment in mice bearing MHC-II^high^ or MHC-II^low^ tumor cells, which corresponded to T cell high or low TMEs by flow cytometry (**Fig. 4H**), respectively. In tumor cells classified as MHC- II^low^, cobimetinib treatment induced a trend toward increased MHC-II expression in malignant PDAC cells in tumor-bearing mice. In contrast, MHC-II^high^ tumor cells showed no appreciable change in MHC-II expression following cobimetinib treatment in vivo (**Fig. 4H–I**). Notably, cobimetinib treatment led to a significant reduction in tumor volume in mice bearing MHC- II^low^ tumor cells, which paralleled the observed upregulation of MHC-II on these same (previously MHC-II^low^) tumor cells after therapy in vitro and in vivo (**Fig. 4 G–J**).

Next, we examined the therapeutic impact of combining cobimetinib in vivo with or without dual immune checkpoint blockade (ICB: both anti-PD-1 and anti-CTLA-4; **Fig. 4K**). Mice bearing tumors and treated with the combination of cobimetinib + ICB therapy exhibited substantially reduced tumor growth compared to vehicle + isotype control treated mice across both MHC-II^high^ and MHC-II^low^ tumor bearing mice, although the most pronounced benefit was observed in mice bearing MHC-II^low^ tumors (**Fig. 4L–M**). Specifically, the MHC- II^low^ tumor clone 6419c5—derived from the same KPC model reported by in Li et al.^25^, and known to be highly therapeutically resistant—responded robustly to the combination of cobimetinib + ICB even in the orthotopic setting (**Fig. 4N**). This therapeutic regimen not only reduced tumor burden but also significantly prolonged survival, indicating a marked therapeutic advantage when MHC-II expression was paired with immunotherapy even in therapy-resistant, MHC-II^low^ tumor clones.

Although MHC-II^high^ tumor clones showed partial MHC-II upregulation after cobimetinib stimulation in vitro (**Fig. 4G**) and did not exhibit further upregulation of MHC-II gene expression following cobimetinib treatment in vivo (**Fig. 4I**), MHC-II^high^ TMEs still responded to cobimetinib + ICB therapy (**Fig. 4L**). Given that PDAC is typically considered an T cell- cold tumor, this finding indicates that baseline MHC-II activity in malignant cells may underlie the variability in patient responses to ICB therapy^60^. Furthermore, in MHC-II^low^ tumors, cobimetinib + ICB significantly reduced tumor volume and improved survival relative to ICB monotherapy in both subcutaneous and orthotopic models (**Fig. 4N**; **Supplementary** Fig.

**10A**). In contrast, this combination therapy yielded minimal benefit over ICB alone in MHC- II^high^ tumors (**Supplementary** Fig. 10B), consistent with the lack of MHC-II expression change in these tumor cells observed following cobimetinib treatment (**Fig. 4H-J**).

Overall, these findings highlight MHC-II expression as a key determinant of immunotherapy responsiveness in PDAC with MHC-II^high^ tumors mounting more effective immune-mediated tumor control, and importantly, increasing MHC-II expression in MHC-II^low^ tumor clones can overcome resistance to immune checkpoint monotherapy. Collectively, these results underscore MHC-II as both a predictive biomarker and a promising therapeutic target for enhancing the efficacy of select immunotherapy in pancreatic cancer.

## Discussion

Pancreatic ductal adenocarcinoma remains one of the deadliest malignancies, characterized by its dismal prognosis and highly T cell-suppressive tumor microenvironment. The complexities associated with PDAC, such as its extensive stromal component and heterogeneous immune landscape^61^, necessitate a deeper understanding of how immune dynamics influence tumor progression and therapeutic response. This study provides crucial insights into the role of tumor-intrinsic MHC class II expression on malignant PDAC cells as a modulator of immune responses and highlights its potential as a predictive biomarker for immunotherapy responsiveness.

Our findings reveal that MHC-II expression by malignant PDAC cells correlates strongly with enhanced CD4⁺ and CD8⁺ T-cell infiltration. This observation underscores the potential of malignant PDAC cells to engage adaptive immune responses through antigen presentation, a role traditionally attributed to professional antigen-presenting cells. Elevated MHC-II expression fosters a more immunologically active TME, characterized by higher densities of effector immune cells and enhanced anti-tumor immunity. This aligns with previous studies in other malignancies, such as melanoma^23^, non-small cell lung cancer (NSCLC)^62^, breast cancer^63^ and colorectal cancer^64^, where tumor-intrinsic MHC-II expression was associated with improved clinical outcomes and increased immunogenicity.

In addition to the promising associations observed, our study also identified a subset of PDAC specimens characterized by high MHC-II expression but minimal lymphocyte infiltration, particularly within datasets generated using single-nucleus RNA sequencing. This discrepancy underscores a key limitation of certain single-cell technologies, which tend to underrepresent immune populations due to their preferential capture of malignant and stromal compartments^47,48^.

In addition, we observed substantial intratumoral heterogeneity in MHC-II expression, with MHC-II^high^ and MHC-II^low^ malignant cell niches coexisting within the same tumor microenvironment—consistently across both human and mouse models. Notably, although scRNA-seq data revealed minimal correlation between plasma cell abundance and MHC-II pathway activity, spatial transcriptomic analysis revealed a pronounced spatial proximity between plasma cells and MHC-II^high^ malignant cell regions. These findings underscore the limitations of dissociative profiling methods that disrupt tissue architecture and fail to capture the spatial and temporal heterogeneity critical for interpreting immune-malignant cell interactions^65^. In contrast, spatial transcriptomics preserves native tissue organization, providing a physiologically relevant view of how localized cellular interactions—such as those between immune and malignant cells—influence tumor behavior and immune modulation. The observed spatial co-localization highlights the unique power of spatially resolved technologies to uncover biologically meaningful relationships that remain inaccessible to conventional single-cell approaches.

Interestingly, our analysis revealed that pro-inflammatory CAF subtypes, specifically apCAFs and iCAFs, are predominantly enriched in MHC-II^high^ malignant cell niches.

Conversely, CAF subtypes associated with tumor progression, such as mCAFs and tCAFs, are more prevalent around MHC-II^low^ malignant cell regions. These spatial patterns were consistent across metastatic sites and remained robust regardless of patient sex and age, indicating that while factors like germline background and environmental exposures may contribute to the observed heterogeneity, they are unlikely to be the principal drivers of TME composition. Amidst the increasingly fragmented and complex molecular classification of tumor subtypes, the framework of cancer hallmarks offers a powerful lens through which to interpret these dynamics^66^. Within this context, MHC-II expression by malignant cells may represent a unifying feature—both reflective of and influential on the T cell-suppressive or immune-active state of distinct tumor niches. By spatially aligning with specific fibroblast populations and immune cell distributions, MHC-II may function as a central axis that integrates diverse ecological cues within the TME, ultimately shaping the trajectory of tumor progression. The ability of MHC-II expression to predict therapeutic outcomes carries significant clinical implications, particularly for patient stratification and the design of effective combination therapies. Incorporating malignant cell MHC-II expression levels into diagnostic workflows could enable the identification of patients more likely to benefit from immune checkpoint inhibitors, thereby guiding personalized treatment strategies.

Among the findings of this study, the most compelling is the strong correlation between malignant cell-intrinsic MHC-II expression and responsiveness to current immunotherapy. In vivo experiments demonstrated that tumors with low baseline MHC-II malignant cell expression were unresponsive to combined chemo- and immunotherapy regimens. However, therapeutic induction of MHC-II expression in these tumors led to a marked reversal of treatment resistance. Mechanistically, the upregulation of MHC-II following MEK inhibition may be attributed to relief of ERK-mediated repression of the MHC-II transcriptional machinery.

Previous studies have shown that hyperactivation of the MAPK/ERK pathway—frequently observed in KRAS-mutant tumors—can suppress the expression of CIITA, the master regulator of MHC-II gene transcription^24,67^. MEK inhibition attenuates ERK signaling, thereby lifting this transcriptional block and promoting CIITA expression. Moreover, building on findings from melanoma study^68^, where IFN-γ enhances antigen presentation and promotes T cell– mediated immune responses, we speculate that MEK inhibition may also sensitize tumor cells to IFN-γ. This enhanced responsiveness could further amplify MHC-II gene induction via the STAT1/IRF1 pathway, contributing to a more immunogenic tumor phenotype. Together, these mechanisms suggest that MEK inhibition not only restores intrinsic MHC-II transcriptional activity but also potentiates extrinsic cytokine-driven immune activation, thereby transforming immunologically “cold” tumors into more immune-permissive niches. Specifically, interventions using cobimetinib or combination with IFN-γ successfully upregulated malignant cell MHC-II levels, enhanced T cell infiltration, and converted immunologically “cold” tumors into immunotherapy-responsive TMEs. These results indicate that augmenting MHC-II expression by malignant PDAC cells may represent a viable strategy to overcome primary resistance to checkpoint inhibitor therapy. This approach holds promise for improving clinical outcomes in PDAC patients, particularly those who would otherwise be refractory to current immunotherapeutic options.

While our study provides compelling evidence supporting the tumor-cell intrinsic role of MHC-II in shaping the immune landscape and influencing checkpoint inhibitor responses, several limitations merit further exploration. First, the observed heterogeneity in MHC-II expression across different datasets underscores the need for standardized methodologies to accurately quantify MHC-II levels in clinical samples. Second, the regulatory mechanisms governing MHC-II expression in malignant PDAC cells remain incompletely understood.

Furthermore, the dynamics of interactions between cytotoxic immune cells and malignant PDAC cells with high MHC-II expression are yet to be fully elucidated. Additionally, the influence of other TME components, such as T cell-suppressive myelomonocytic cells, neutrophils, and regulatory T cells (Tregs), on MHC-II-mediated immune responses warrants further investigation. Understanding the complex interplay between MHC-II-expressing malignant PDAC cells and both immunostimulatory and immunosuppressive populations could shed light on how tumors are eliminated or evade immune surveillance despite high MHC-II expression. Lastly, another limitation is the inability to precisely track status changes of individual tumor cells on-treatment in vivo, despite efforts to maintain consistent sampling positions when collecting matched single-cell data. This technological constraint introduces potential bias in our analysis. However, the robust findings from our in vivo mouse experiments reinforce the significant role of MHC-II expression levels in influencing treatment efficacy.

Future research will focus on uncovering the underlying drivers of variability in MHC-II expression among malignant PDAC cells. Specifically, we aim to investigate whether this heterogeneity is driven by intrinsic factors, such as genetic mutations like KRAS and tumor evolution, or by extrinsic cues within the TME. Gaining deeper insights into these regulatory mechanisms will be crucial for developing more precise and effective therapeutic strategies.

In summary, our findings position malignant PDAC cell MHC-II expression as a pivotal modulator of the immune landscape in PDAC and highlight its dual role as a prognostic biomarker and a potential therapeutic target. By promoting antigen presentation and fostering an immune-supportive TME, MHC-II expression enhances the efficacy of immunotherapy and represents a promising avenue for therapeutic intervention. Strategies that upregulate MHC-II expression in malignant PDAC cells, such as IFN-γ stimulation or combination treatments targeting the MAPK pathway, could help overcome therapeutic resistance and improve patient outcomes in this highly lethal disease. Future clinical trials are warranted to evaluate the safety and efficacy of MHC-II-targeted therapies and to further clarify their potential to transform the treatment landscape for PDAC.

## Methods

### Human tissue samples

This study was conducted in accordance with the ethical principles outlined in the Helsinki Declaration. The dataset comprised 2 PDAC tissue samples from clinical patient ST00022941, including matched pre- and on-treatment biopsies from a patient enrolled in a window-of-opportunity SMMART trial involving cobimetinib therapy (NCT04005690).

Collection and handling of human patient-derived materials adhered to protocols approved by the Oregon Health & Science University (OHSU) Institutional Review Board (IRB #00003609). All tissue samples were obtained from the Oregon Pancreas Tissue Registry program. Written informed consent was obtained from all patients prior to sample collection. Patients were not compensated for participation.

### Tissue dissociation

Patient tumor tissues were obtained from surgical resections and dissociated using gentleMACS Dissociator kits (Miltenyi Biotec, #130-095-929) following the manufacturer’s instructions. The resulting lysates were resuspended and filtered through a 70-μm cell strainer (Miltenyi Biotec, #130-098-462). Cells were collected by centrifugation at 300 × g for 7 minutes at 4°C and subsequently resuspended at a concentration of 700–1,200 cells/μL. Live cells were enriched using the EasySep™ Dead Cell Removal (Annexin V) Kit (STEMCELL Technologies, #17899).

### Single-Cell RNA Sequencing Library Construction

Single-cell suspensions were processed according to the 10x Genomics single-cell RNA-seq protocol using the Chromium Next GEM Single Cell 3’ Kit v3.1. No prior sorting or enrichment was performed, allowing analysis of the entire mixed cell population. Cell suspensions were loaded onto the Chromium Controller to generate Gel Beads-in-Emulsion (GEMs). Libraries were prepared from amplified cDNA using 14 PCR cycles. Quality control of intermediate products and final libraries was performed using a 1% E-Gel system (Thermo Fisher Scientific), and library concentrations were quantified using a Qubit Fluorometer (Thermo Fisher Scientific), following the manufacturer’s protocols. All libraries from a given patient were prepared within the same experimental batch to minimize batch effects. Sequencing was carried out on the Illumina NovaSeq 6000 platform at the OHSU Massively Parallel Sequencing Shared Resource, with a targeted sequencing depth of 20,000 read pairs per cell.

### scRNA-seq data processing and quality control

Raw sequencing data were processed using Cell Ranger (10x Genomics, v7.1.0). The mkfastq and count commands were executed with default parameters, aligning reads to the GRCh38 human reference genome to generate a matrix of unique molecular identifier (UMI) counts per gene and corresponding cell barcodes. Seurat objects were constructed from each dataset using the Seurat pipeline (version 5.0.3).

Quality control filters were applied to exclude low-quality cells and potential doublets.

Specifically, cells were removed if they met any of the following criteria: fewer than 300 detected genes or fewer than 1,200 UMIs; greater than 20% mitochondrial gene expression, indicating compromised cell integrity; greater than 7,500 UMI counts and more than 60,000 transcripts, suggestive of doublets. The filtered gene expression matrix was normalized using the NormalizeData function with default parameters. The merged and normalized matrix was used for downstream analysis. Doublets were computationally identified and removed using DoubletFinder (version 2.0.4) with the following parameters: pN (proportion of artificial doublets): 0.25 (default value); nExP (expected doublet rate): 0.075 (based on the DoubletFinder pipeline). These settings were provided as inputs to the doubletFinder function, with the number of principal components (PCs) set to 10 for doublet identification.

Cells were clustered using the Louvain algorithm and the FindNeighbors and FindClusters functions, with the top 10 PCA dimensions and a resolution parameter of 0.5. The Uniform Manifold Approximation and Projection (UMAP) for visualizing clusters was generated using the RunUMAP function with the same 10 principal components used for clustering.

### scRNA-seq cell-type annotation

Cell-type annotation was performed by assigning main cell types to each cluster based on the expression of a comprehensive set of established marker genes. The expression profiles were manually reviewed to ensure accuracy and consistency. All cell-type assignments were performed by a single annotator to maximize consistency across all datasets.

### Mice

C57BL/6 (Cat# 00664) mice were purchased from Jackson Laboratories (Bar Harbor, ME) and were bred and maintained under specific pathogen-free conditions at an American Association for the Accreditation of Laboratory Animal Care (AAALAC)-accredited animal facilities within Oregon Health and Science University. Mice were housed in compliance with the procedures that were reviewed and approved by the Institutional Animal Care and Use Committee of Oregon Health and Science University (Protocol #4153). Unless otherwise stated, sex and age-matched littermates (6-12 weeks of age, both sexes) were used for individual experiments.

### *In vivo* reagents and treatments

Mice were treated with cobimetinib (Selleck Chemicals; Cat# S8041) by oral gavage at a dose of 5.0 mg/kg, in 10% PEG300/PBS. Treatment commenced after detectable tumors; between 7-14 days for orthotopically implanted tumors or 10-12 days for subcutaneously implanted tumors. Immune checkpoint blockade (‘ICB’) was treated by dual treatment with αPD- 1/αCTLA-4 (‘PC’) as previously described^69^. Briefly, αPD-1 (RMP1-14; BioXcell, Cat # BE0146; 200 μg/dose) and αCTLA-4 (9H10; BioXcell, Cat # BE0131; 200 μg/dose) were injected intraperitoneally (i.p.) on days 0, 3, and 6^70^. For isotype controls, rat IgG2a (2A3; BioXcell, Cat # BE0089; 200 μg) was used. All antibodies were endotoxin free.

### Implantation of tumor cell clones

Kras^LSL-G12D/+^; Trp53^LSL-R172H/+^; Pdx1-Cre; Rosa26^YFP/YFP^ tumor clones were generated as previously described^25^ and are also commercially available (Kerafast). Cells were cultured in DMEM (high glucose without sodium pyruvate) with 7.5-10% FBS (Gibco) and glutamine (2.0 mM) and harvested when confluent. Cells were dissociated into single cells with 0.25% trypsin (Gibco), washed with serum-free Dulbecco’s Modified Eagle’s medium (DMEM) twice, and counted in preparation for subcutaneous implantation at a dose of 2.5x10^5^ PDAC tumor cells. For orthotopic tumors, 0.5-5x10^5^ PDAC cells were implanted into the pancreas via ultrasound guided injection^71^ and monitored by ultrasound every 5-7 days. Tumor cells had viability >90% for each experiment, and cell lines were examined by the Infectious Microbe PCR Amplification Test (IMPACT) and authenticated to be free of contamination by the Research Animal Diagnostic Laboratory (RADIL) at the University of Missouri.

### Tumor growth, regression, and animal survival assessment

Subcutaneous tumor sizes were measured every 2-3 days for tumor growth assessment experiments. Tumor length and width were measured with calipers and tumor volumes were then calculated as length*width^2^/2. Orthotopic tumors were detected and monitored longitudinally by ultrasound every 5-7 days using a Fujifilm VevoSonic 2100 ultrasound with 3D motor. Tumor regressions and waterfall plots were calculated using the initial tumor size at the start of treatment to tumor size at indicated later time point.

### Flow cytometry of murine tumor samples

At the indicated time points, mice were sacrificed and tumors prepared for single cell suspension as previously described^70^. Briefly, tumors were washed with PBS, then minced and incubated for 45 minutes in 1mg/mL collagenase XI with protease inhibitor (Sigma-Aldrich, Cat # C6079) at 37°C and filtered through a 70 μM cell strainer with cold PBS supplemented with 0.5% BSA and 2.0 mM ethylenediaminetetraacetic acid (EDTA). Cells were counted with Beckman Coulter Counter Z2 and stained with Ghost Dye UV450 (Cytek Biosciences Cat # 13-0868-T100). Cell surface molecules were assessed by incubating single cell suspensions from tissues with primary fluorophore-conjugated antibodies on ice for 45 minutes in PBS with 0.5% bovine serum albumin and 2.0 mM EDTA. Combinations of the following antibodies were used for flow cytometric analysis: I-A/I-E (MHC-II) (clone M5/114.15.2, Brilliant Violet 711, BD Horizons, Cat # 563414), CD3e (clone 145-2C11, PE-Cy5, Biolegend, Cat # 100310), CD45 (clone 30-F11, UV486, BD Optibuild, Cat # 749889). Flow cytometric analysis was performed on Fortessa flow cytometer (BD Biosciences) using BD FACSDiva software and analyzed using FlowJo software (Treestar).

For in vitro tumor cell stimulations, the IC50 for each cell line after in vitro treatment with cobimetinib was determined by MTT Assay. Tumor cell lines were then cultured +/- cobimetinib at IC50 (0.0004-0.5 μg/mL) or vehicle control, +/- 10 ng/mL recombinant mouse IFN-γ, for 48 hours. Tumor cell expression of I-A (MHC-II) was then determined by flow cytometry as above.

### Multiplexed Immunohistochemistry

Tumor specimen slides were processed and stained with a panel of 22 antibodies following previously described protocols^42,72^. Multiple 1.0 mm^2^ regions of interest (ROIs) were sampled across each primary tumor resections based on tissue quality after staining representative of the whole tissue sample. Regions were annotated using Aperio Image Scope (Leica Biosystems). Image registration was performed using the SURF algorithm in the Computer Vision Toolbox from MATLAB (The MathWorks), and the AEC chromogen signal was extracted using Fiji^73^. Single-cell segmentation and labeling were carried out with StarDist 2D, and marker mean signal intensities were quantified at the single-cell level using CellProfiler^74^. Gating thresholds for marker positivity were established using FCS Express Image Cytometry (De Novo Software).

### Single-cell data collection and processing

Public scRNA-seq data were sourced from the Deeply Integrated Single-Cell Omics (DISCO) database (https://www.immunesinglecell.org/). A total of 22 datasets comprising 264 pancreatic samples and 1,042,407 cells were collected to build the PDAC atlas. This included 115 PDAC samples encompassing 557,304 cells, which were used to investigate the relationship between MHC-II pathway activity in malignant cells or predicted responsive malignant cell proportions and the infiltration of other cell types. To address potential batch effects arising from different platforms and protocols, the BBKNN integration algorithm was applied with default parameters. Clustering and single-cell distributions were visualized using UMAP with the Leiden clustering algorithm. Malignant cells were identified using the R package scATOMIC (v2.0.2)^45^, with the parameter known_cancer_type = "PAAD cell" and default settings for all other parameters. Spearman’s correlation analysis was conducted to evaluate associations between MHC-II pathway activity or the proportion of responsive malignant cells and cell-type infiltration in each sample.

### Spatial transcriptomic data collection and processing

Public spatial transcriptomic RNA sequencing datasets were obtained from four sources. First, 14 Visium samples and their matched snRNA-seq data from the WUSTL Atlas were downloaded from the Human Tumor Atlas Network (HTAN) Data Coordinating Center Data Portal at the National Cancer Institute (https://data.humantumoratlas.org/). Second, 55 Visium samples, encompassing both primary and metastatic lesions, were retrieved from the Gene Expression Omnibus (GEO) under accession number GSE277783. As this second set lacked matched single-cell transcriptomic data, we utilized the same publicly available scRNA-seq dataset^75^ employed in Pei’s study^37^, derived from 23 patients with PDAC. This dataset contained 23,042 cells and captures all major cell populations, including T cells, B cells, plasma cells, myeloid cells, fibroblasts, endothelial cells, and malignant tumor cells.

Additionally, four well-annotated Visium samples and nine ST sample were obtained from the SORC database (http://bio-bigdata.hrbmu.edu.cn/SORC/Browse.jsp#). Lastly, two Xenium slices from human pancreatic cancer tissue—generated using the pre-designed Xenium Human Multi-Tissue and Cancer Panel—were downloaded from the 10x Genomics database. For downstream analysis, cells from each Xenium dataset were filtered to retain those with nFeature_Xenium > 10 and nCount_Xenium > 20.

The RCTD method was used to deconvolute cell-type composition within each spatial spot, utilizing scRNA-seq data as the reference for the 10x Visium samples from the HTAN WUSTL cohort. RCTD objects were generated from processed RDS files with the parameters max_cores = 20 and CELL_MIN_INSTANCE = 10. The RCTD pipeline was executed on these objects with doublet_mode = "full" for 10x Visium data. The deconvolution results were stored as matrices of cell-type weights for each spot. The cell-type weights for each spot were normalized to ensure that the sum of weights equaled 1. Proportion matrices were created, with rows representing spatial spots and columns representing cell types and were stored for further analysis. For rough annotation, each spatial spot was assigned a primary cell type based on the highest probabilistic proportion.

### Bulk dataset collection and preprocessing

In this study, RNA-seq fastq data for 39 sorted mouse tumor cell samples (PRJNA432498) were downloaded from the GEO database (http://www.ncbi.nlm.nih.gov/geo/). The quality of the raw fastq files was assessed using FastQC (v0.11.8) (http://www.bioinformatics.bbsrc.ac.uk/projects/fastqc/). Sequencing reads were aligned to the GRCm38 mouse reference genome using STAR (v2.6.0a), and per-gene counts and transcripts per kilobase million (TPM) values were quantified using RSEM (v1.3.1) based on the gencode.vM25.annotation.gtf gene annotation file. For normalization, gene lengths were determined by the number of bases covered at least once across all exons for each gene.

TPM values were calculated as follows: 1. Reads per kilobase (RPK) were computed for each gene; 2. The scaling factor was calculated as the sum of all RPK values Scaling factor = ∑_!_ RPK_!_; 3. TPM values were derived by normalizing each gene’s RPK to the scaling factor and multiplying by 10^6^: 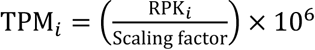. For cases with duplicate sequencing runs for the same patient, TPM counts were averaged. Gene expression values were log-transformed using the formula log_3_(TPM + 1) for each sample. To standardize expression data, z-scores were calculated for each gene across all samples using the formula: 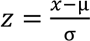 where *x* is the log-transformed TPM value, µ is the average log_3_(TPM + 1) across all samples for a given gene, and σ is the standard deviation of log_3_(TPM + 1) for that gene.

### Copy-number analysis

Copy-number variations in malignant epithelial cells were inferred using InferCNV (version 1.18.1). Lymphoid immune cells, including CD8⁺ T cells, CD4⁺ T cells, NK cells, and B cells, were used as reference controls. The input data consisted of gene expression count matrices. Filtering, normalization, and centering based on normal gene expression were performed using the default parameters, followed by data scaling. A minimum average read count threshold of 0.1 per gene among reference cells was applied to ensure robust CNV detection.

### Differential scRNA gene expression and pathway enrichment analysis

Differential gene expression analysis was performed using two-tailed Wilcoxon rank-sum tests implemented in the Seurat package via the FindMarkers function with default parameters. Results were visualized using dot plots to depict the relative expression levels of differentially expressed genes (DEGs) across experimental conditions. Gene Ontology (GO) analysis was performed using the msigdbr R package (v7.5.1). Rank-based gene set enrichment analysis (GSEA) was conducted using the clusterProfiler R package (v4.10.0). For bulk transcriptomic data and pseudo-bulk data (generated by aggregating gene expression counts from all malignant PDAC cells using the PseudobulkExpression module in the Seurat package), single-sample GSEA (ssGSEA) was performed with the GSVA R package (v1.50.0). For scRNA-seq and stRNA-seq data, the AddModuleScore function in Seurat was utilized to account for technical variability and mitigate the impact of dropouts.

Genes of interest were binned into 24 expression-level bins based on their average expression. Control genes were randomly selected from each bin using default parameters in the AddModuleScore function. The final module score was calculated as the difference between the average expression of the gene set of interest and the average expression of the control genes.

### Determining cancer-cell lineage states

To characterize the heterogeneity of cancer cell lineage states, we assessed MHC-II expression across multiple platforms. For Visium data, the AddModuleScore function was used to calculate MHC-II expression scores. Spots with a score greater than 0 were classified as “MHC-II^high^,” while those with a score of 0 or below were designated “MHC-II^low^”. For Xenium data, due to the sparsity of gene expression signals, MHC-II status in cancer cells was defined based on the detectable expression of canonical MHC-II markers. For mIHC data, MHC-II protein expression status in malignant cells were determined using sample-specific thresholds adjusted for background noise, ensuring that marker positivity accurately reflected true expression.

### Cellular neighborhood analyses

To characterize the spatial relationships between immune, stromal, and tumor cells, cellular neighborhoods were defined by placing a circular window with a fixed radius of 50 µm around each seed cell of a specified phenotype. All cells whose centers fell within this radius were considered neighbors and were included in the corresponding seed cell’s neighborhood. Neighborhoods across all malignant cell regions were then analyzed by clustering their normalized cellular compositions. This was performed using the dbscan R package in combination with the k-nearest neighbors (kNN) algorithm, followed by K-means clustering to identify distinct spatial neighborhood types. To assess compositional differences among the neighborhoods of malignant cells, we applied the Wilcoxon rank-sum test. Both absolute cell counts and relative proportions of neighboring cell types were evaluated to determine statistically significant variations. To ensure robust and reliable comparisons in the downstream cellular neighborhood analyses, each Visium sample was required to meet the following criteria: a minimum of 10 spots classified as “MHC-II^high^” and 10 spots as “MHC-II^low^”, as well as either at least 5 immune-enriched spots for immune neighborhood analyses or a minimum of 50 CAF spots for stromal neighborhood analyses, respectively.

### Heterotypic interaction between spatially adjacent cell types

To quantify spatial interactions between distinct cell types, we computed a heterotypic interaction score based on spatial proximity within the Visium platform. First, spatial coordinates of each spot were scaled and mapped to a unified spatial object. For each sample, the six nearest spatial neighbors of each spot were identified using the kNN function like before, with a maximum distance threshold of 200 µm and a minimum of four ranked neighbors retained to ensure local connectivity. The resulting neighbor network was converted into an undirected graph containing weighted edges inversely proportional to spatial distances between spots. To examine heterotypic interactions, we utilized cell type deconvolution matrices obtained from previous analyses, focusing on spot pairs enriched for different cell types. A spot was considered positive for a given cell type if its deconvolution score exceeded 0.2. For each pair of distinct cell types, we identified all edges in the spatial network that connected spots enriched for one cell type to those enriched for another. These edges were used to define the heterotypic interaction network between the two cell types.

### Statistical Analysis

All statistical analyses were conducted using R (v4.3.1) and GraphPad Prism (versions 7). For comparisons between two independent groups, either a two-tailed unpaired Student’s t-test (for normally distributed data) or Wilcoxon rank-sum test (for non-normally distributed data) was applied. Kaplan–Meier survival analyses were assessed using the log-rank test. Tumor growth kinetics were analyzed using two-way ANOVA with mixed effects modeling. For analyses involving multiple comparisons, the Benjamini–Hochberg procedure was used to control the false discovery rate. Error bars in all figures represent either the standard deviation (SD) or the standard error of the mean (SEM), as indicated in the figure legends. Statistical significance was defined as *p* < 0.05. Significance levels are denoted as follows: *p* < 0.05 (*), *p* < 0.01 (**), and *p* < 0.001 (***). “ns” indicates not significant.

## Acknowledgments

This work was supported by the following funding sources: the Department of Defense Prostate Cancer Data Science Award (HT94252410551), NIH R01GM147365, and the Silver Family Innovation Foundation Award (to Z.X.); NIH P30 CA069533, NIH U01CA253472, U01CA281902 and U24CA264128 (to G.B.M.); The Susan G. Komen Foundation, the National Foundation for Cancer Research and Hildegard Lamfrom Endowed Chair in Basic Research, and NIH P30 CA069533 (to L.M.C.); NIH U01CA294548 and U01CA278923 (to R.C.S.) and the Robert L. Fine Cancer Research Foundation and the Knight Cancer Institute (K.T.B.). The SMMART clinical trial and sample collections, L.M.C, R.C.S, G.B.M. and K.T.B. were supported by the Brenden-Colson Center for Pancreatic Care and by the Oregon Pancreas Tissue Registry. Targeted Pathway Inhibition in Patients with Pancreatic Cancer ClinicalTrials.gov ID NCT04005690. We would like to acknowledge the Human Tumor Atlas Network (HTAN) for providing valuable data (phs002371.v1.p1). The content is solely the responsibility of the authors and does not necessarily represent the official views of the funders.

## Author contributions

Z.X. conceived the idea. C.C. performed the data analysis. J.K., Y.Z., K.G., and K.T.B. conducted the experimental validations. X.L., T.O., and F.O. generated scRNA-seq data. S.S. and K.E.B. performed mIHC data generation and analysis. X.Y. and R.S.D. contributed to data analysis. M.H.S. and M.S.D. contributed to result interpretation. D.K., R.C.S., and C.D.L. coordinated the clinical trial and provided clinical insights. L.M.C. led the mIHC analysis and interpreted results. G.B.M., K.T.B. and Z.X. supervised the study. C.C., K.T.B., and Z.X. wrote the manuscript. All other authors edited the manuscript and provided critical feedback and approved the final manuscript.

## Data availability

Publicly available datasets used in this study include the Xenium spatial transcriptomics datasets from 10x Genomics, and Visium samples from the HTAN WUSTL atlas, which are accessible via the Human Tumor Atlas Network (HTAN) dbGaP Study Accession: phs002371.v1.p1 (https://www.ncbi.nlm.nih.gov/projects/gap/cgi-bin/study.cgi?study_id=phs002371.v1.p1), as well as through the Gene Expression Omnibus (GEO) under accession number GSE277783. Additional spatial data were obtained from the SORC^39^ database. Cancer cell line expression data were retrieved from the Cancer Cell Line Encyclopedia (https://sites.broadinstitute.org/ccle/datasets). The human genome reference (GRCh38, version 4.0.0) used for single-cell analyses was obtained from public sources, with genome builds and annotation files provided by 10x Genomics (https://support.10xgenomics.com/single-cell-gene-expression/software/release-notes/build). For mouse transcriptomic analyses, the GRCm38 reference genome and GENCODE vM25 annotation were downloaded from the GENCODE portal (https://www.gencodegenes.org/mouse/). The mIHC-derived single-cell phenotyping and spatial localization data have been deposited in Synapse and are accessible at https://www.synapse.org/#!Synapse:syn51078766.

## Code availability

All packages used in this study are publicly available. All data, scripts and code used for preprocessing, statistical analysis, and figure generation are available upon request.

## Supplemental material

Figure S1-S10.

**Supplemental table 1.** MHC-II Gene Expression Levels Across PDAC and Pan-Cancer Tumor Cell Lines.

**Supplemental table 2.** List of scRNA-seq Cohorts Used in This Study.

## Reference

1. Siegel, R. L., Miller, K. D., Wagle, N. S. & Jemal, A. Cancer statistics, 2023. CA: A Cancer J. Clin. 73, 17–48 (2023).

2. Sarantis, P., Koustas, E., Papadimitropoulou, A., Papavassiliou, A. G. & Karamouzis, M. V. Pancreatic ductal adenocarcinoma: Treatment hurdles, tumor microenvironment and immunotherapy. World J. Gastrointest. Oncol. 12, 173–181 (2020).

3. Neoptolemos, J. P. et al. Therapeutic developments in pancreatic cancer: current and future perspectives. Nat. Rev. Gastroenterol. Hepatol. 15, 333–348 (2018).

4. Makohon-Moore, A. & Iacobuzio-Donahue, C. A. Pancreatic cancer biology and genetics from an evolutionary perspective. Nat. Rev. Cancer 16, 553–565 (2016).

5. Gamboa, A. C. et al. Optimal timing and treatment strategy for pancreatic cancer. J. Surg. Oncol. 122, 457–468 (2020).

6. Rahib, L., Fleshman, J. M., Matrisian, L. M. & Berlin, J. D. Evaluation of Pancreatic Cancer Clinical Trials and Benchmarks for Clinically Meaningful Future Trials: A Systematic Review. JAMA Oncol. 2, 1209 (2016).

7. Zhang, J., Späth, S. S., Marjani, S. L., Zhang, W. & Pan, X. Characterization of cancer genomic heterogeneity by next-generation sequencing advances precision medicine in cancer treatment. *Precis*. Clin. Med. 1, 29–48 (2018).

8. Hirata, E. & Sahai, E. Tumor Microenvironment and Differential Responses to Therapy. Cold Spring Harb. Perspect. Med. 7, a026781 (2017).

9. Li, X. et al. Panoramic tumor microenvironment in pancreatic ductal adenocarcinoma. Cell. Oncol. 47, 1561–1578 (2024).

10. Peng, J. et al. Single-cell RNA-seq highlights intra-tumoral heterogeneity and malignant progression in pancreatic ductal adenocarcinoma. Cell Res. 29, 725–738 (2019).

11. Shalek, A. K. & Benson, M. Single-cell analyses to tailor treatments. Sci. Transl. Med. 9, eaan4730 (2017).

12. Tellez-Gabriel, M., Ory, B., Lamoureux, F., Heymann, M.-F. & Heymann, D. Tumour Heterogeneity: The Key Advantages of Single-Cell Analysis. Int. J. Mol. Sci. 17, 2142 (2016).

13. Dai, Z. et al. Research and application of single-cell sequencing in tumor heterogeneity and drug resistance of circulating tumor cells. Biomark. Res. 8, 60 (2020).

14. Herting, C. J., Karpovsky, I. & Lesinski, G. B. The tumor microenvironment in pancreatic ductal adenocarcinoma: current perspectives and future directions. Cancer Metastasis Rev. 40, 675–689 (2021).

15. Liu, X. et al. The reciprocal regulation between host tissue and immune cells in pancreatic ductal adenocarcinoma: new insights and therapeutic implications. Mol. Cancer 18, 184 (2019).

16. Clark, C. E. et al. Dynamics of the Immune Reaction to Pancreatic Cancer from Inception to Invasion. Cancer Res. 67, 9518–9527 (2007).

17. Ino, Y. et al. Immune cell infiltration as an indicator of the immune microenvironment of pancreatic cancer. Br. J. Cancer 108, 914–923 (2013).

18. Goulart, M. R., Stasinos, K., Fincham, R. E. A., Delvecchio, F. R. & Kocher, H. M. T cells in pancreatic cancer stroma. World J. Gastroenterol. 27, 7956–7968 (2021).

19. Liu, R., Liao, Y.-Z., Zhang, W. & Zhou, H.-H. Relevance of Immune Infiltration and Clinical Outcomes in Pancreatic Ductal Adenocarcinoma Subtypes. Front. Oncol. 10, 575264 (2021).

20. Winograd, R. et al. Induction of T-cell Immunity Overcomes Complete Resistance to PD-1 and CTLA-4 Blockade and Improves Survival in Pancreatic Carcinoma. Cancer Immunol. Res. 3, 399–411 (2015).

21. Amhis, N., Carignan, J. & Tai, L.-H. Transforming pancreaticobiliary cancer treatment: Exploring the frontiers of adoptive cell therapy and cancer vaccines. Mol. Ther.: Oncol. 32, 200825 (2024).

22. Roche, P. A. & Furuta, K. The ins and outs of MHC class II-mediated antigen processing and presentation. Nat. Rev. Immunol. 15, 203–216 (2015).

23. Axelrod, M. L., Cook, R. S., Johnson, D. B. & Balko, J. M. Biological Consequences of MHC-II Expression by Tumor Cells in Cancer. Clin. Cancer Res. 25, 2392–2402 (2019).

24. Johnson, A. M. et al. Cancer Cell–Intrinsic Expression of MHC Class II Regulates the Immune Microenvironment and Response to Anti–PD-1 Therapy in Lung Adenocarcinoma. J. Immunol. 204, 2295–2307 (2020).

25. Li, J. et al. Tumor Cell-Intrinsic Factors Underlie Heterogeneity of Immune Cell Infiltration and Response to Immunotherapy. Immunity 49, 178–193.e7 (2018).

26. Marum, L. Cancer Cell Line Encyclopedia launched by Novartis and Broad Institute. *Futur*. Med. Chem. 4, 947 (2012).

27. Hu, C. et al. CellMarker 2.0: an updated database of manually curated cell markers in human/mouse and web tools based on scRNA-seq data. Nucleic Acids Res. 51, D870–D876 (2022).

28. Zhou, D. C. et al. Spatially restricted drivers and transitional cell populations cooperate with the microenvironment in untreated and chemo-resistant pancreatic cancer. Nat. Genet. 54, 1390–1405 (2022).

29. inferCNV of the Trinity CTAT Project. https://github.com/broadinstitute/inferCNV.

30. Oketch, D. J. A., Giulietti, M. & Piva, F. Copy Number Variations in Pancreatic Cancer: From Biological Significance to Clinical Utility. Int. J. Mol. Sci. 25, 391 (2023).

31. Sakamoto, H. et al. The Evolutionary Origins of Recurrent Pancreatic Cancer. Cancer Discov. 10, 792–805 (2020).

32. Yu, G., Wang, L.-G., Han, Y. & He, Q.-Y. clusterProfiler: an R Package for Comparing Biological Themes Among Gene Clusters. *OMICS: A J*. Integr. Biol. 16, 284–287 (2012).

33. Yi, R. et al. MHC-II Signature Correlates With Anti-Tumor Immunity and Predicts anti-PD-L1 Response of Bladder Cancer. Front. Cell Dev. Biol. 10, 757137 (2022).

34. Liu, C. et al. Single-cell RNA-sequencing reveals radiochemotherapy-induced innate immune activation and MHC-II upregulation in cervical cancer. Signal Transduct. Target. Ther. 8, 44 (2023).

35. Yabo, Y. A., Niclou, S. P. & Golebiewska, A. Cancer cell heterogeneity and plasticity: A paradigm shift in glioblastoma. Neuro-Oncol. 24, 669–682 (2021).

36. Bhat, G. R. et al. Cancer cell plasticity: from cellular, molecular, and genetic mechanisms to tumor heterogeneity and drug resistance. Cancer Metastasis Rev. 43, 197–228 (2024).

37. Pei, G. et al. Spatial mapping of transcriptomic plasticity in metastatic pancreatic cancer. Nature 1–10 (2025) doi:10.1038/s41586-025-08927-x.

38. Cable, D. M. et al. Robust decomposition of cell type mixtures in spatial transcriptomics. Nat. Biotechnol. 40, 517–526 (2022).

39. Zhou, W. et al. SORC: an integrated spatial omics resource in cancer. Nucleic Acids Res. 52, D1429–D1437 (2023).

40. 10x Genomics Xenium 1.6.0.8 using 10x Genomics Cloud Analysis.

41. Banik, G. et al. High-dimensional multiplexed immunohistochemical characterization of immune contexture in human cancers. Methods Enzym. 635, 1– 20 (2019).

42. Liudahl, S. M. et al. Leukocyte Heterogeneity in Pancreatic Ductal Adenocarcinoma: Phenotypic and Spatial Features Associated with Clinical Outcome. Cancer Discov. 11, 2014–2031 (2021).

43. Li, M. et al. DISCO: a database of Deeply Integrated human Single-Cell Omics data. Nucleic Acids Res. 50, D596–D602 (2021).

44. Polañski, K. et al. BBKNN: fast batch alignment of single cell transcriptomes. Bioinformatics 36, 964–965 (2019).

45. Nofech-Mozes, I., Soave, D., Awadalla, P. & Abelson, S. Pan-cancer classification of single cells in the tumour microenvironment. Nat. Commun. 14, 1615 (2023).

46. Hwang, W. L. et al. Single-nucleus and spatial transcriptome profiling of pancreatic cancer identifies multicellular dynamics associated with neoadjuvant treatment. Nat. Genet. 54, 1178–1191 (2022).

47. Klughammer, J. et al. A multi-modal single-cell and spatial expression map of metastatic breast cancer biopsies across clinicopathological features. Nat. Med. 30, 3236–3249 (2024).

48. Slyper, M. et al. A single-cell and single-nucleus RNA-Seq toolbox for fresh and frozen human tumors. Nat. Med. 26, 792–802 (2020).

49. Zeng, Z. et al. Immune and stromal scoring system associated with tumor microenvironment and prognosis: a gene-based multi-cancer analysis. J. Transl. Med. 19, 330 (2021).

50. Dakal, T. C., George, N., Xu, C., Suravajhala, P. & Kumar, A. Predictive and Prognostic Relevance of Tumor-Infiltrating Immune Cells: Tailoring Personalized Treatments against Different Cancer Types. Cancers 16, 1626 (2024).

51. Bruni, D., Angell, H. K. & Galon, J. The immune contexture and Immunoscore in cancer prognosis and therapeutic efficacy. Nat. Rev. Cancer 20, 662–680 (2020).

52. Mirlekar, B. et al. Balance between immunoregulatory B cells and plasma cells drives pancreatic tumor immunity. Cell Rep. Med. 3, 100744 (2022).

53. Wei, R. et al. Spatial charting of single-cell transcriptomes in tissues. Nat. Biotechnol. 40, 1190–1199 (2022).

54. Cords, L. et al. Cancer-associated fibroblast classification in single-cell and spatial proteomics data. Nat. Commun. 14, 4294 (2023).

55. Joshi, R. S. et al. The Role of Cancer-Associated Fibroblasts in Tumor Progression. Cancers 13, 1399 (2021).

56. Wright, K., Ly, T., Kriet, M., Czirok, A. & Thomas, S. M. Cancer-Associated Fibroblasts: Master Tumor Microenvironment Modifiers. Cancers 15, 1899 (2023).

57. Huang, H. et al. Mesothelial cell-derived antigen-presenting cancer-associated fibroblasts induce expansion of regulatory T cells in pancreatic cancer. Cancer Cell 40, 656–673.e7 (2022).

58. Zarmer, S., McCabe, I. & Yeh, J. J. Abstract 494: Identifying vulnerabilities in basal-like pancreatic adenocarcinoma (PDAC). Cancer Res. 85, 494–494 (2025).

59. Hingorani, S. R. et al. Trp53R172H and KrasG12D cooperate to promote chromosomal instability and widely metastatic pancreatic ductal adenocarcinoma in mice. Cancer Cell 7, 469–483 (2005).

60. Bear, A. S., Vonderheide, R. H. & O’Hara, M. H. Challenges and Opportunities for Pancreatic Cancer Immunotherapy. Cancer Cell 38, 788–802 (2020).

61. Wang, S. et al. The molecular biology of pancreatic adenocarcinoma: translational challenges and clinical perspectives. Signal Transduct. Target. Ther. 6, 249 (2021).

62. Johnson, A. M. et al. Cancer Cell-Specific Major Histocompatibility Complex II Expression as a Determinant of the Immune Infiltrate Organization and Function in the NSCLC Tumor Microenvironment. J. Thorac. Oncol. 16, 1694–1704 (2021).

63. Gonzalez-Ericsson, P. I. et al. Tumor-Specific Major Histocompatibility-II Expression Predicts Benefit to Anti–PD-1/L1 Therapy in Patients With HER2- Negative Primary Breast Cancer. Clin. Cancer Res. 27, 5299–5306 (2021).

64. Warabi, M., Kitagawa, M. & Hirokawa, K. Loss of MHC class II expression is associated with a decrease of tumor-infiltrating T cells and an increase of metastatic potential of colorectal cancer Immunohistological and histopathological analyses as compared with normal colonic mucosa and adenomas. Pathol. - Res. Pr. 196, 807– 815 (2000).

65. Desta, G. M. & Birhanu, A. G. Advancements in single-cell RNA sequencing and spatial transcriptomics: transforming biomedical research. Acta Biochim. Pol. 72, 13922 (2025).

66. Sibai, M. et al. The spatial landscape of cancer hallmarks reveals patterns of tumor ecological dynamics and drug sensitivity. Cell Rep. 44, 115229 (2025).

67. Yao, Y. et al. ERK and p38 MAPK Signaling Pathways Negatively Regulate CIITA Gene Expression in Dendritic Cells and Macrophages. J. Immunol. 177, 70–76 (2006).

68. Wawrzyniak, P. & Hartman, M. L. Dual role of interferon-gamma in the response of melanoma patients to immunotherapy with immune checkpoint inhibitors. Mol. Cancer 24, 89 (2025).

69. Morrison, A. H., Diamond, M. S., Hay, C. A., Byrne, K. T. & Vonderheide, R. H. Sufficiency of CD40 activation and immune checkpoint blockade for T cell priming and tumor immunity. Proc. Natl. Acad. Sci. 117, 8022–8031 (2020).

70. Morrison, A. H., Diamond, M. S., Hay, C. A., Byrne, K. T. & Vonderheide, R. H. Sufficiency of CD40 activation and immune checkpoint blockade for T cell priming and tumor immunity. Proc National Acad Sci 201918971 (2020) doi:10.1073/pnas.1918971117.

71. Hay, C. A., Sor, R., Flowers, A. J., Clendenin, C. & Byrne, K. T. Ultrasound- Guided Orthotopic Implantation of Murine Pancreatic Ductal Adenocarcinoma. J Vis Exp (2019) doi:10.3791/60497.

72. Link, J. M. et al. Ongoing replication stress tolerance and clonal T cell responses distinguish liver and lung recurrence and outcomes in pancreatic cancer. *Nat*. Cancer 6, 123–144 (2025).

73. Schindelin, J., et al. Fiji: an open-source platform for biological-image analysis. Nat. Methods 9, 676–682 (2012).

74. Carpenter, A. E. et al. CellProfiler: image analysis software for identifying and quantifying cell phenotypes. Genome Biol. 7, R100 (2006).

75. Raghavan, S. et al. Microenvironment drives cell state, plasticity, and drug response in pancreatic cancer. Cell 184, 6119–6137.e26 (2021).

