## Supplementary Figures for "Tumor-intrinsic MHC-II activation in pancreatic ductal adenocarcinoma enhances immune response and treatment efficacy"

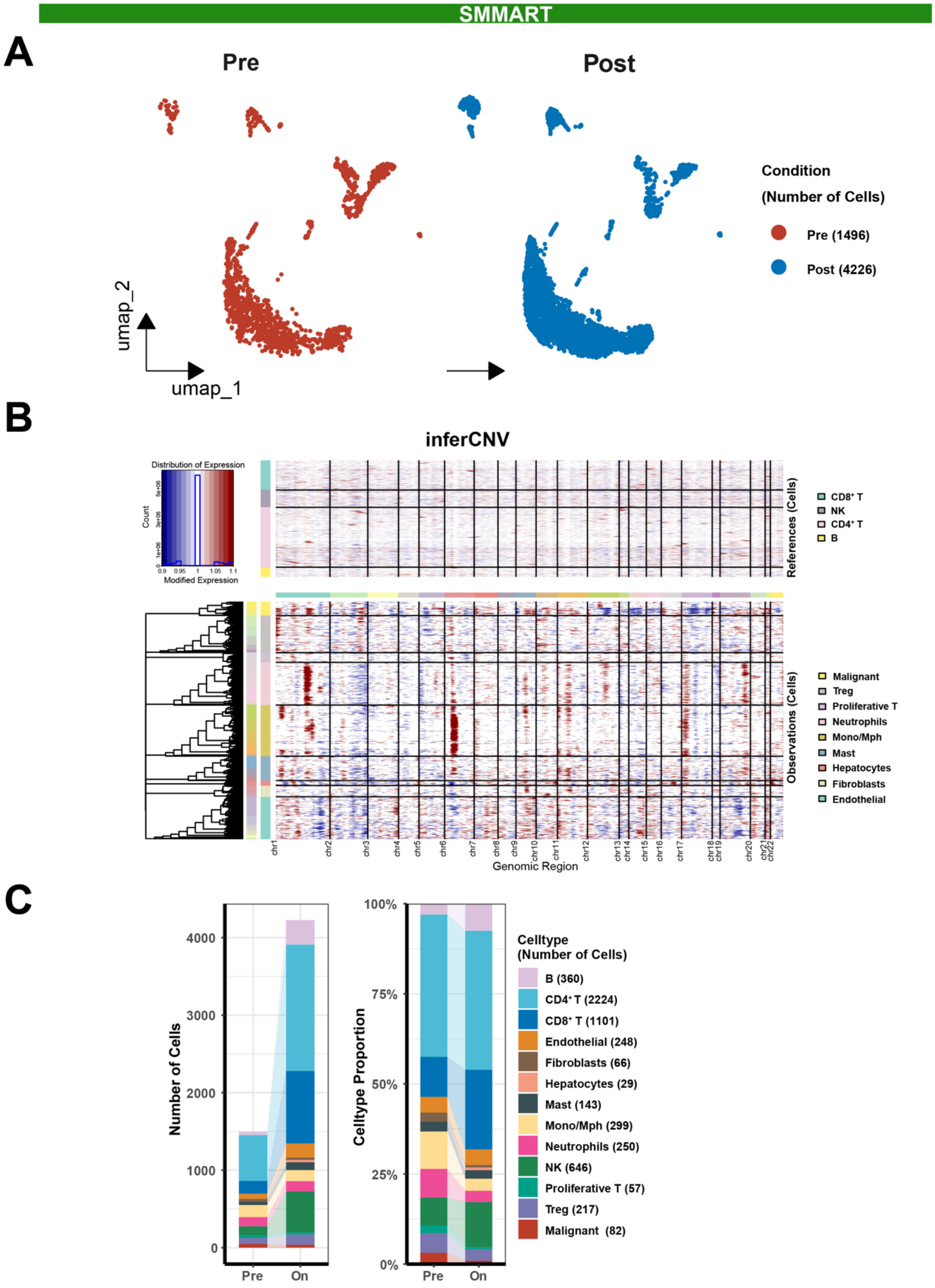

SMMART
A
Pre
Post
Condition
(Number of Cells)
Pre (1496)
Post (4226)
umap_2
umap_1
B
inferCNV
Distribution of Expression
Count
Modified Expression
References (Cells)
CD8⁺ T
NK
CD4⁺ T
B
Observations (Cells)
Malignant
Treg
Proliferative T
Neutrophils
Mono/Mph
Mast
Hepatocytes
Fibroblasts
Endothelial
chr1 chr2 chr3 chr4 chr5 chr6 chr7 chr8 chr9 chr10 chr11 chr12 chr13 chr14 chr15 chr16 chr17 chr18 chr19 chr20 chr21 chr22
Genomic Region
C
Number of Cells
4000
3000
2000
1000
0
Pre
On
Celltype Proportion
100%
75%
50%
25%
0%
Celltype
(Number of Cells)
B (360)
CD4⁺ T (2224)
CD8⁺ T (1101)
Endothelial (248)
Fibroblasts (66)
Hepatocytes (29)
Mast (143)
Mono/Mph (299)
Neutrophils (250)
NK (646)
Proliferative T (57)
Treg (217)
Malignant (82)

**Supplementary Figure 1. Pre- and on-treatment scRNA-seq analysis of liver metastases in PDAC patients undergoing cobimetinib therapy. A.** UMAP plot showing clustering of pre- (red) and on-treatment (blue) single-cell profiles. Minimal batch effects are observed, indicating robust integration of datasets across conditions. Cell numbers are indicated in parentheses for each condition. **B.** Copy number variation analysis of single-cell gene expression. Chromosomal alterations are evident in malignant epithelial cells, confirming their malignant status. Reference cells include $CD8^+$ T cells, NK cells, $CD4^+$ T cells, and B cells. **C.** Bar plots representing the number of cells (left) and cell-type proportions (right) for pre- and on-treatment samples.

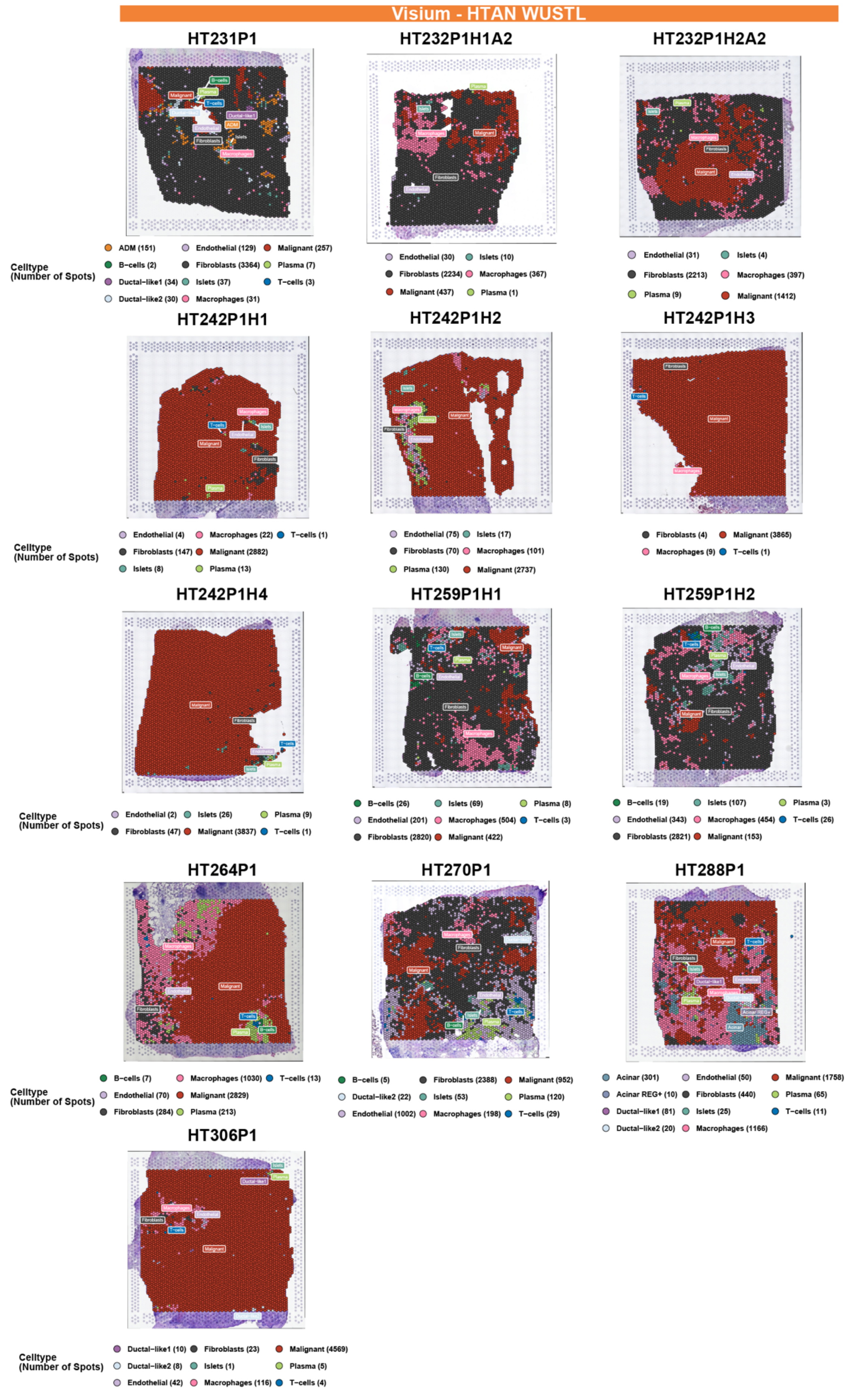

Visium - HTAN WUSTL
HT231P1
ADM (151) Endothelial (129) Malignant (257)
B−cells (2) Fibroblasts (3364) Plasma (7)
Ductal−like1 (34) Islets (37) T−cells (3)
Ductal−like2 (30) Macrophages (31)
Celltype (Number of Spots)
HT232P1H1A2
Endothelial (30) Islets (10)
Fibroblasts (2234) Macrophages (367)
Malignant (437) Plasma (1)
HT232P1H2A2
Endothelial (31) Islets (4)
Fibroblasts (2213) Macrophages (397)
Plasma (9) Malignant (1412)
HT242P1H1
Celltype (Number of Spots)
Endothelial (4) Macrophages (22) T−cells (1)
Fibroblasts (147) Malignant (2882)
Islets (8) Plasma (13)
HT242P1H2
Endothelial (75) Islets (17)
Fibroblasts (70) Macrophages (101)
Plasma (130) Malignant (2737)
HT242P1H3
Fibroblasts (4) Malignant (3865)
Macrophages (9) T−cells (1)
HT242P1H4
Celltype (Number of Spots)
Endothelial (2) Islets (26) Plasma (9)
Fibroblasts (47) Malignant (3837) T−cells (1)
HT259P1H1
B−cells (26) Islets (69) Plasma (8)
Endothelial (201) Macrophages (504) T−cells (3)
Fibroblasts (2820) Malignant (422)
HT259P1H2
B−cells (19) Islets (107) Plasma (3)
Endothelial (343) Macrophages (454) T−cells (26)
Fibroblasts (2821) Malignant (153)
HT264P1
Celltype (Number of Spots)
B−cells (7) Macrophages (1030) T−cells (13)
Endothelial (70) Malignant (2829)
Fibroblasts (284) Plasma (213)
HT270P1
B−cells (5) Fibroblasts (2388) Malignant (952)
Ductal−like2 (22) Islets (53) Plasma (120)
Endothelial (1002) Macrophages (198) T−cells (29)
HT288P1
Acinar (301) Endothelial (50) Malignant (1758)
Acinar REG+ (10) Fibroblasts (440) Plasma (65)
Ductal−like1 (81) Islets (25) T−cells (11)
Ductal−like2 (20) Macrophages (1166)
HT306P1
Celltype (Number of Spots)
Ductal−like1 (10) Fibroblasts (23) Malignant (4569)
Ductal−like2 (8) Islets (1) Plasma (5)
Endothelial (42) Macrophages (116) T−cells (4)

**Supplementary Figure 2. Spatial transcriptomics analysis of PDAC tumor regions depicting cell type distributions.** Spatial transcriptomics maps for PDAC samples generated using Visium data from the HTAN WUSTL atlas. Each map illustrates the distribution of distinct cell types within tumor regions, and cell counts per type are indicated in the legend.

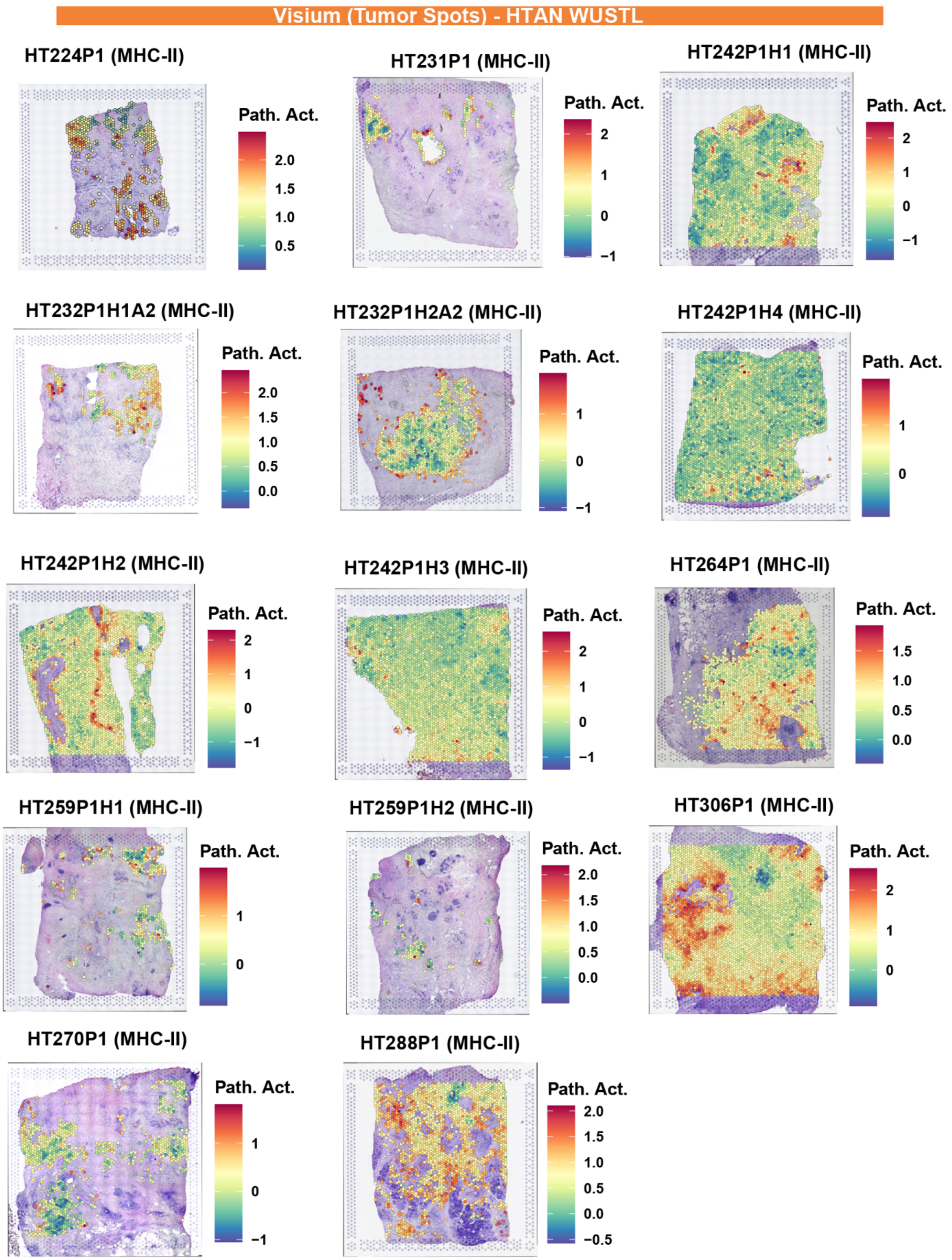
Visium (Tumor Spots) - HTAN WUSTL
HT224P1 (MHC-II)
Path. Act.
2.0
1.5
1.0
0.5
HT231P1 (MHC-II)
Path. Act.
2
1
0
−1
HT242P1H1 (MHC-II)
Path. Act.
2
1
0
−1
HT232P1H1A2 (MHC-II)
Path. Act.
2.0
1.5
1.0
0.5
0.0
HT232P1H2A2 (MHC-II)
Path. Act.
1
0
−1
HT242P1H4 (MHC-II)
Path. Act.
1
0
HT242P1H2 (MHC-II)
Path. Act.
2
1
0
−1
HT242P1H3 (MHC-II)
Path. Act.
2
1
0
−1
HT264P1 (MHC-II)
Path. Act.
1.5
1.0
0.5
0.0
HT259P1H1 (MHC-II)
Path. Act.
1
0
HT259P1H2 (MHC-II)
Path. Act.
2.0
1.5
1.0
0.5
0.0
HT306P1 (MHC-II)
Path. Act.
2
1
0
HT270P1 (MHC-II)
Path. Act.
1
0
−1
HT288P1 (MHC-II)
Path. Act.
2.0
1.5
1.0
0.5
0.0
−0.5

**Supplementary Figure 3. Spatial transcriptomics analysis of MHC-II pathway activity in PDAC malignant cell spots.** Spatial feature plots representing the overall MHC-II pathway activity across rest PDAC tumor samples. Each plot includes a color scale denoting normalized expression or pathway activity levels.

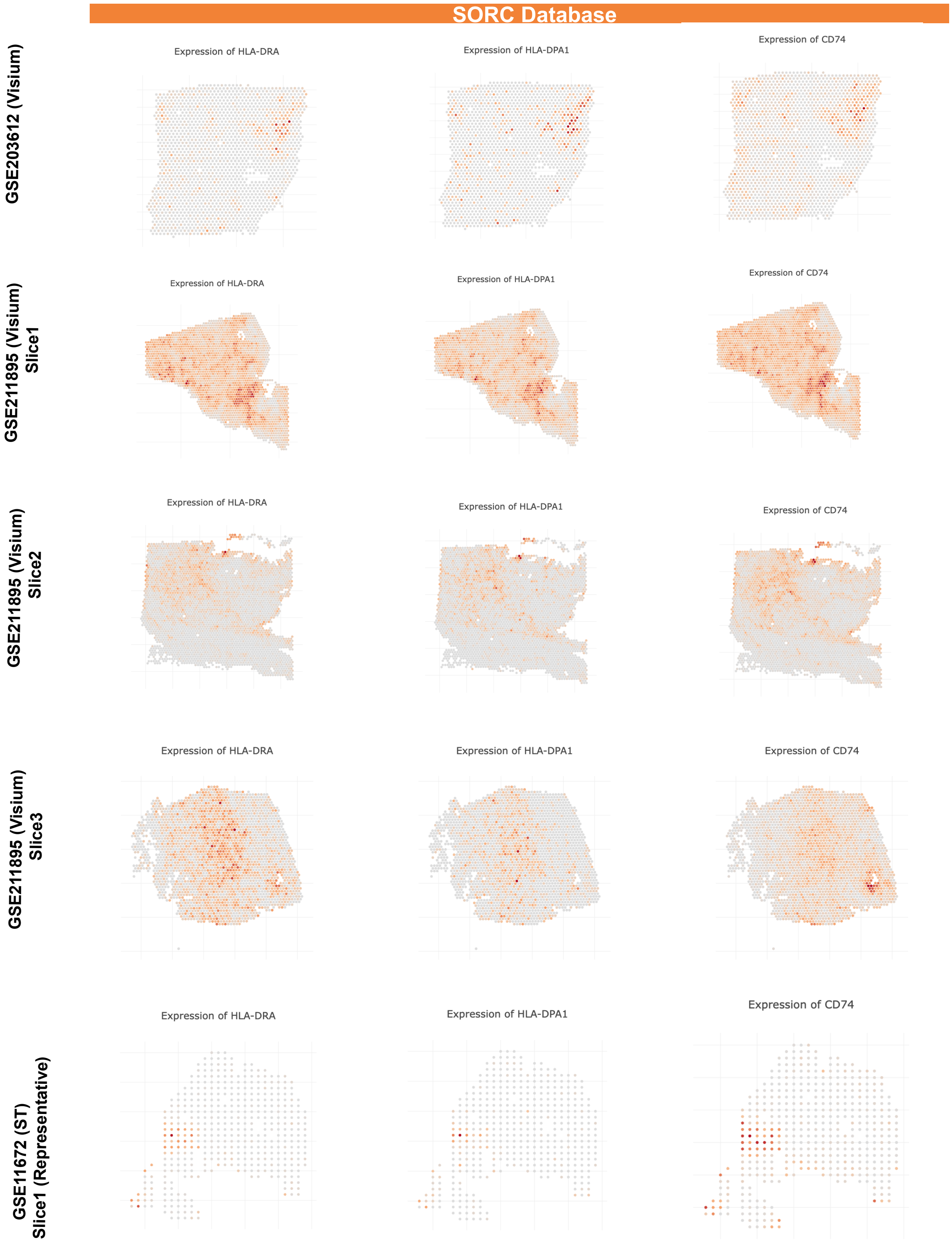
SORC Database
GSE203612 (Visium)
GSE211895 (Visium) Slice1
GSE211895 (Visium) Slice2
GSE211895 (Visium) Slice3
GSE11672 (ST) Slice1 (Representative)
Expression of HLA-DRA
Expression of HLA-DPA1
Expression of CD74

**Supplementary Figure 4. Validation of MHC-II pathway gene expression using spatial transcriptomics from the SORC database.** Spatial expression maps of MHC-II pathway-related genes (*HLA-DRA*, *HLA-DPA1*, and *CD74*) across multiple Visium spatial transcriptomics datasets from the SORC database. Each panel depicts the spatial distribution of normalized expression levels for these genes within representative PDAC tumor slices. High expression regions (red) highlight localized areas of active antigen presentation machinery, while lower expression areas (gray) indicate spatial heterogeneity. The datasets include slices from GSE203612, GSE211895 (Slices 1–3), and a representative slide from GSE211672.

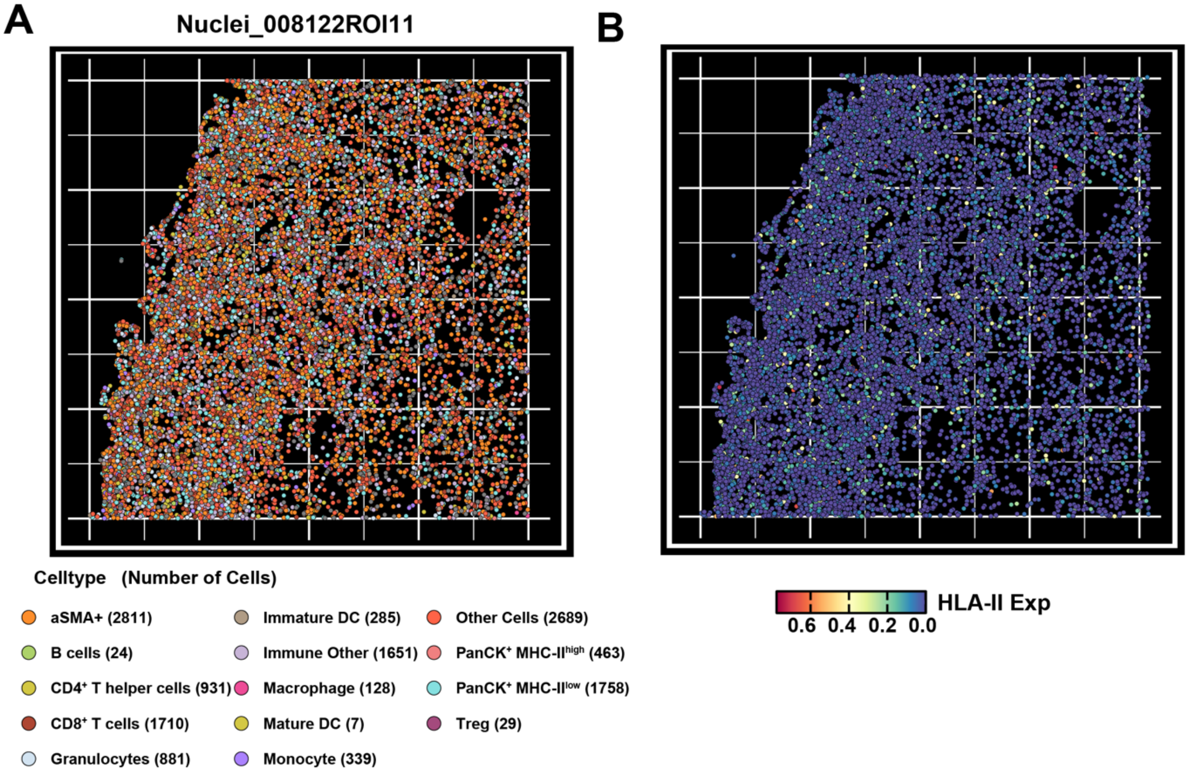


**Supplementary Figure 5. Spatial distribution and HLA-II expression of cell types in a representative mIHC PDAC sample. A.** Multiplex immunohistochemistry (mIHC) image showing the spatial mapping of diverse immune, stromal, and epithelial cell types within region of interest ROI. Cell counts per type are indicated in the legend. **B.** Spatial feature-plot of HLA-II protein expression across the same tissue section.

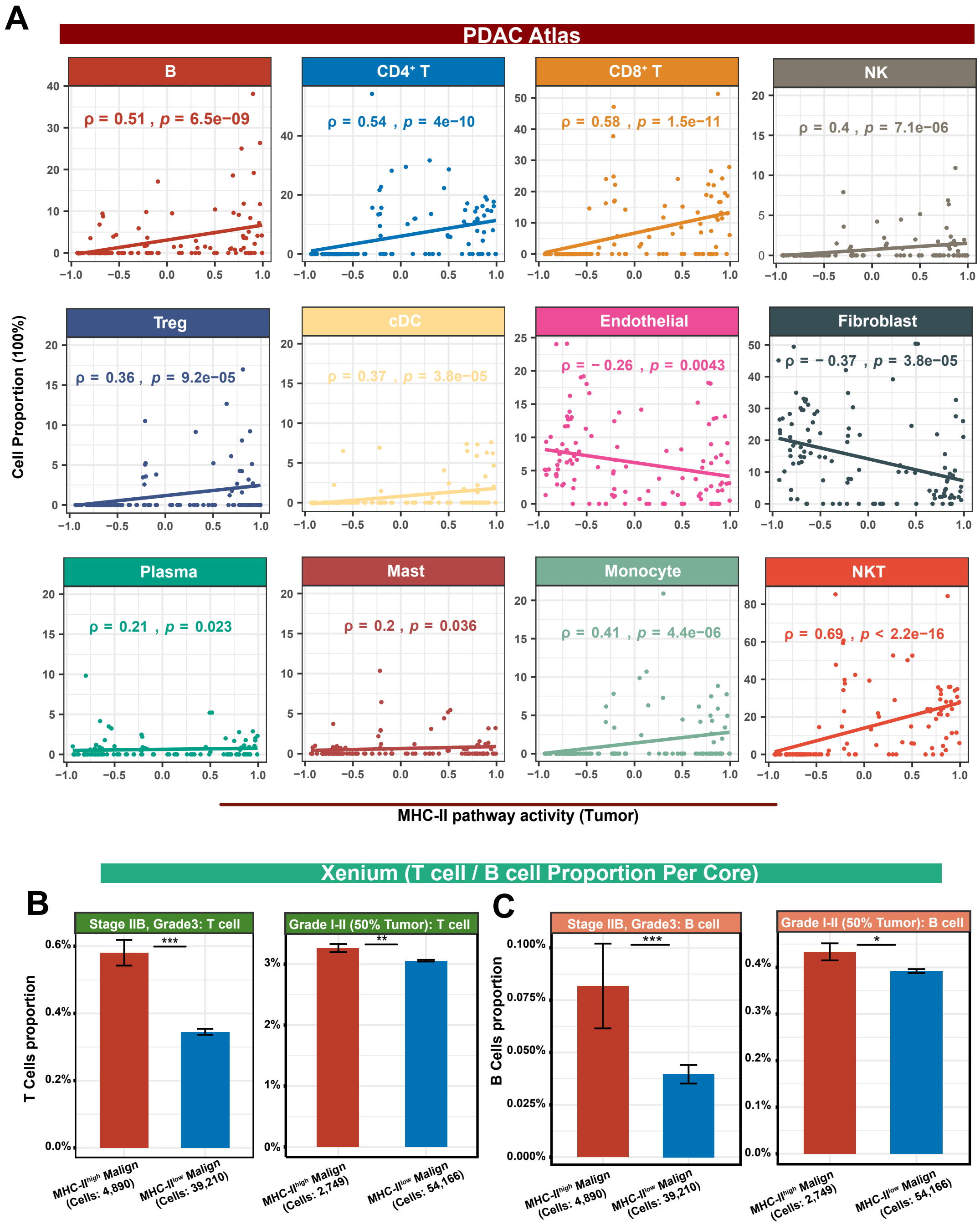


**Supplementary Figure 6. Correlation between MHC-II pathway activity in malignant cells and immune cell proportions in the tumor microenvironment. A.** Scatter plots depicting the Spearman correlation between MHC-II pathway activity in malignant cells and the proportions of various immune and stromal cell types across multiple PDAC samples. Each point represents a sample, with cell proportions expressed as percentages. **B-C.** Bar plots display the proportion of T cells **(B)** and B cells **(C)** within malignant neighborhoods classified as MHC-II$^{high}$ or MHC-II$^{low}$, for two tumor samples: Stage IIB, Grade 3 (left panels) and Grade I–II with ~50% malignant cell content (right panels). Error bars represent standard error of the mean (SEM), and cell counts are indicated for each group. *: $p < 0.05$, **: $p < 0.01$ and ***: $p < 0.001$. The same significance thresholds apply to all subsequent figures unless otherwise specified.

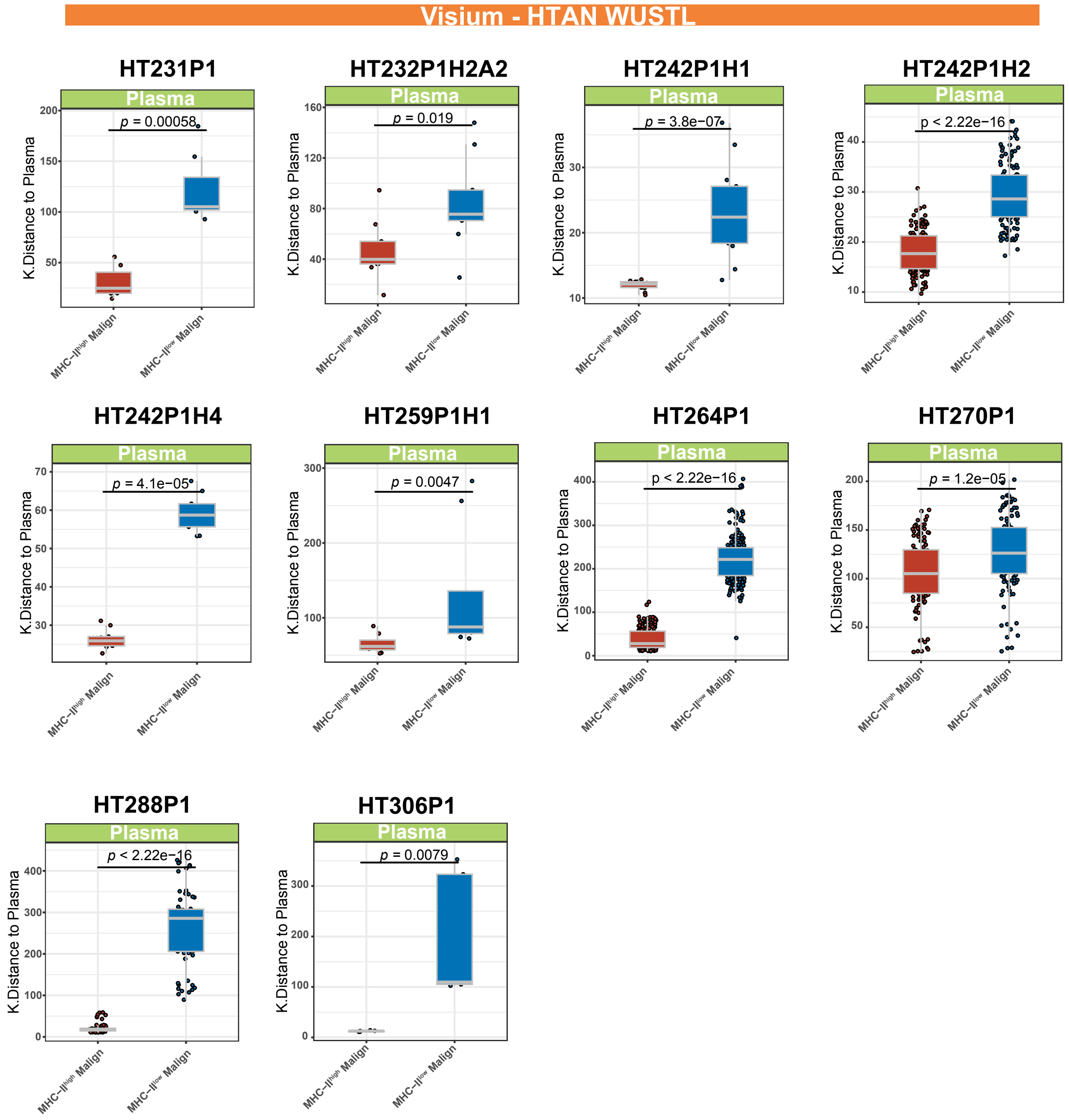


**Supplementary Figure 7. Boxplots display the K-nearest-neighbor (KNN) distances between MHC-II$^{high}$ versus MHC-II$^{low}$ malignant cell spots and plasma cell-enriched regions across spatial transcriptomic samples from the HTAN WUSTL cohort.** Each panel corresponds to an individual slide sample, with statistically significant reductions in closer spatial association between plasma cells and MHC-II$^{high}$ malignant cell spots observed in every case. *P*-values were computed using the Wilcoxon rank-sum test.

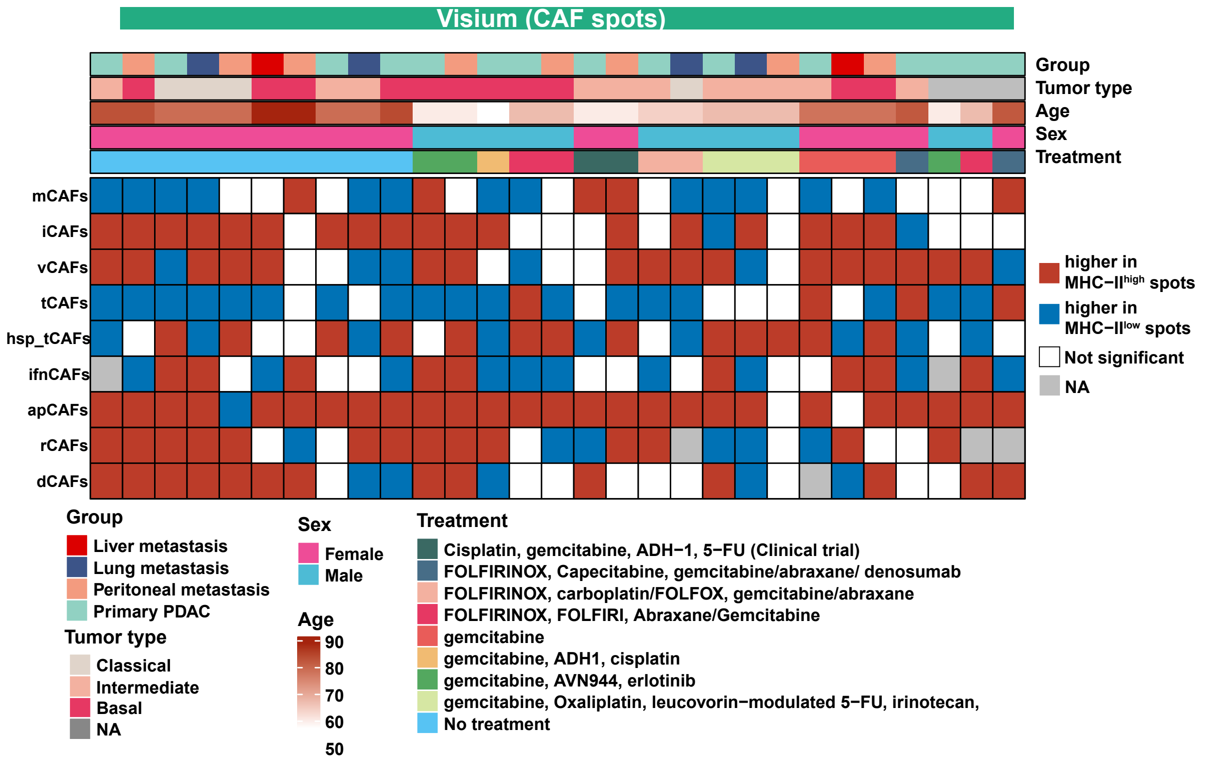


**Supplementary Figure 8. CAF subtype distribution varies by malignant cells' MHC-II pathway activity across Visium samples.** Heatmap summarizing the relative enrichment of nine CAF subtypes across malignant cell spots stratified by MHC-II pathway activity (MHC-II$^{high}$ vs. MHC-II$^{low}$) in spatial transcriptomics data. Each column represents a Visium sample, annotated by metastatic site (group), PDAC tumor type, age, sex, and treatment regimen. Each row corresponds to a specific CAF subtype, including myofibroblastic (mCAFs), inflammatory (iCAFs), vascular (vCAFs), tumor-adjacent (tCAFs), heat shock–associated (hsp_tCAFs), interferon-responsive (ifnCAFs), antigen-presenting (apCAFs), regenerative (rCAFs), and developmental (dCAFs) CAFs.

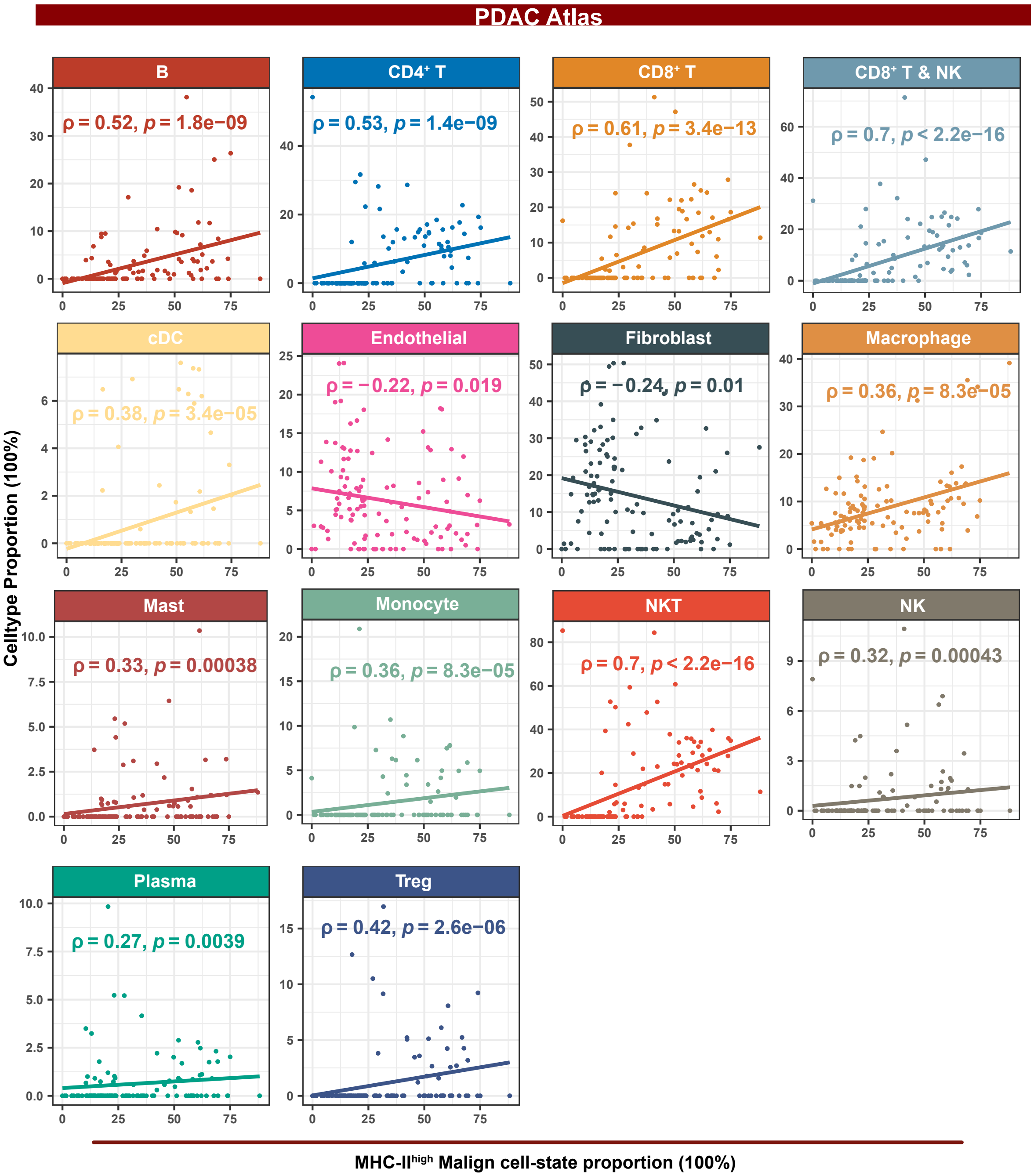


**Supplementary Figure 9. Correlation between the proportion of predicted MHC-II$^{high}$ malignant cells and immune/stromal cell proportions in the PDAC microenvironment.** Scatter plots showing Spearman correlation coefficients (ρ) between the proportion of MHC-II$^{high}$ malignant cells and various cell populations in PDAC samples. Each point represents a PDAC sample, with cell proportions expressed as percentages.

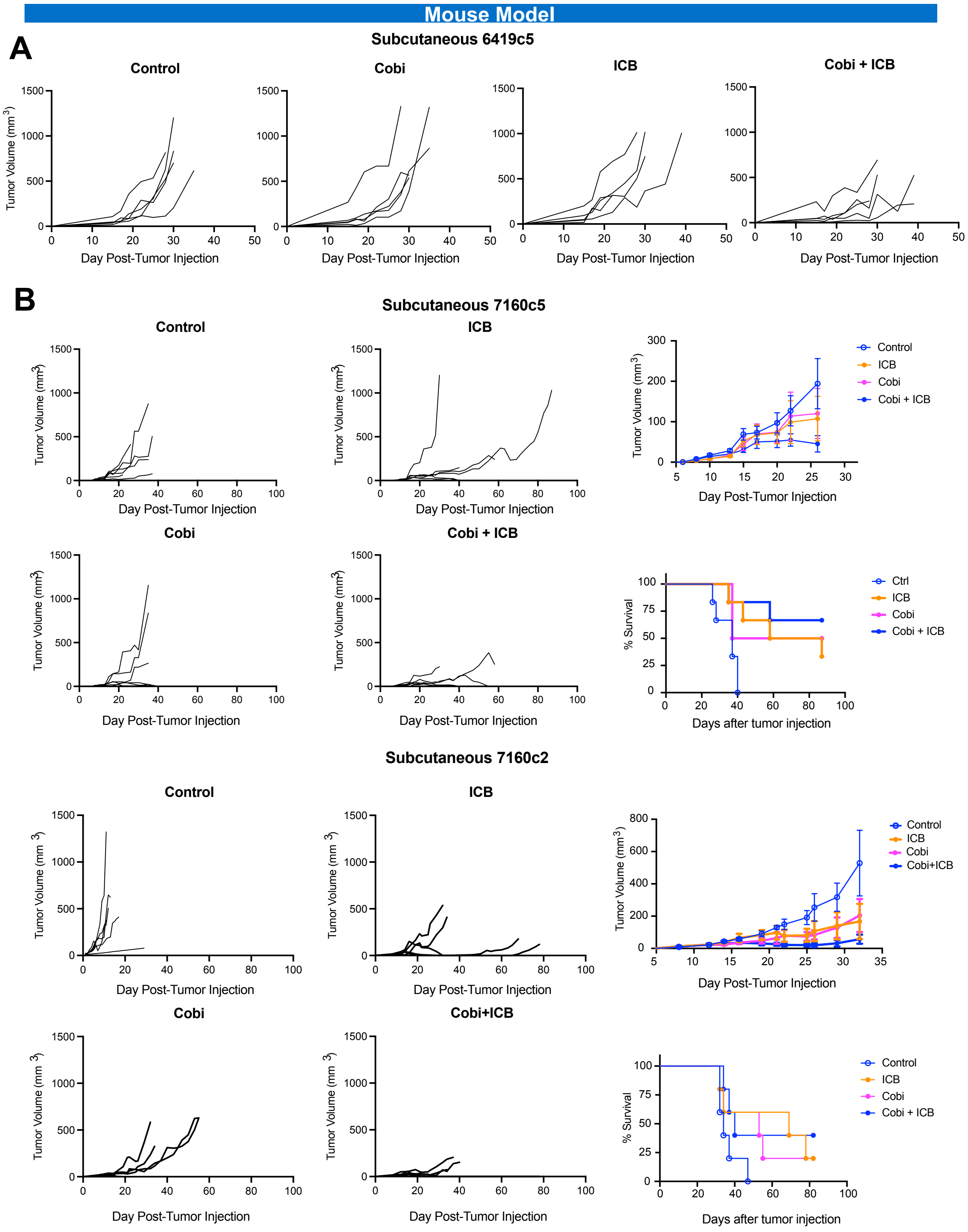


**Supplementary Figure 10. Combination of MEK inhibition and immune checkpoint blockade reduces tumor growth and improves survival in PDAC mouse models. A.**

Individual tumor growth trajectories in subcutaneous 6419c5 tumors (MHC-II$^{low}$) treated with vehicle (Control), cobimetinib (Cobi), immune checkpoint blockade (ICB), or the combination (Cobi + ICB). **B.** Tumor growth and survival outcomes across two MHC-II$^{high}$ subcutaneous PDAC models (7160c5 and 7160c2).
